# AI predicted TCR-pMHC structures differentiate immune interactions

**DOI:** 10.64898/2026.02.24.707744

**Authors:** Michael Robben

## Abstract

The T Cell Receptor (TCR) is a highly variable component of the T cell immune response that recognizes unique epitopes presented on MHC molecules (pMHC). Random genetic recombination limits the ability for sequence homology to predict epitope specificity, which is more dependent on the strength of the TCR-pMHC binding interaction. Structures for understanding this interaction only exist for well characterized positive interactors, and there is no information available about the physical interaction of non-specific TCR-pMHC’s. In this study, we explore the ability for structural prediction algorithms to generate interacting and non-interacting multimeric TCR-pMHC structures, then, examine features that can predict immune interaction. AlphaFold2 shows more consistent multimeric structure prediction compared to other deep learning structure generators or template based algorithms. Poor structure generation does not correlate with immune interaction, and non-interacting structures show similar structural properties to interacting structures. However, this results in less energetically stable conformations in non-interacting structures. Molecular dynamic simulation supports this finding and reveals a novel structural conformation that contributes mechanistically to proper immune synapse. We show that structural and physical features extracted from generated structures are more predictive of interaction than sequence based features. To support researchers in the prediction of TCR-epitope specificity we have made our structural prediction models available through an accessible notebook based webserver: https://github.com/RobbenLab/TCRSIP.

## Introduction

T cells mediate epitope specific immune responses through recognition of foreign and self-antigens presented on the surface of major histocompatibility complex (MHC) molecules^1–3^. The T cell receptor (TCR), which varies per cell due to recombination, recognizes unique 9-13 amino acid peptides and catalyzes internal signaling leading to an immune response. The hypervariable CDR1, CDR2, and CDR3 regions of the TCR interact with the pMHC through predominantly electrostatic interactions and non-covalent bonds.

The relatively low observed binding strength (K_d_ 0.01 – 100 µM)^4^ of the TCR-pMHC interaction and promiscuity of TCRs to multiple epitopes has led to disagreement for the true mechanism responsible for antigen specific responses^5,6^. Rules for specificity have been proposed involving anchor residues, optimal binding angles, and CDR-peptide-MHC interaction but increased structure sampling continuously finds exceptions and high variability^7,8^. Because the TCR does not contain an intracellular signaling region, recent hypotheses for specificity focus on the dynamic events catalyzed during immune synapse. These include kinetic proofreading^9^, mass effect^10^, force-mediated dynamics^11^, and catch-slip bonds^12–14^. Without a clear mode of action, it has been impossible to develop sequence based rules for specificity.

The majority of sequence based predictors focus on the CDR3ab region and peptide sequence because they are the most variable component of synapse^15^. Unsupervised clustering methods are largely no more accurate than classification based on sequence hamming distance^16^. Deep learning algorithms like NetTCR2.0 and ERGO have been found unable to achieve greater than 60-70% accuracy on specificity predictions^17^. These approaches also show a high degree of bias towards the training dataset and inability to generalize to unseen peptides^18^.

New generative AI models such as AlphaFold2^19^, present an opportunity to produce more structural data for TCR-pMHC research. Previously, Bradley^20^ and Yin et al.,^21^ investigated AlphaFold for its use in TCR-pMHC structural prediction. Both used a strategy of optimizing existing AlphaFold 2 models to produce higher fidelity predictions fine-tuned on TCR-pMHC specific structures. Deleuran and Nielsen^22^ proposed further refinement of structural prediction accuracy using a GNN regressor on DOCKQ scores for ranking AlphaFold predicted structures. The strategy utilized by these previous approaches relies on the presumption that non-interacting structures would produce poorly folded structures. In this study, explore how structure prediction algorithms generate non-existing structures and examine the features that are predictive of interaction. We identify physical and biochemical features that can be used to differentiate interacting and non-interacting predicted structures. More interestingly, these features are important for protein interaction as determined by molecular dynamic simulation. Finally, we develop models that can be used to differentiate interacting from non-interacting sequences based on their folded structures.

## Methods and Materials

### Ground truth database preparation

Ground truth examples of TCR-pMHC interacting structures were compiled from the TCR3D database^23^ (downloaded October 14, 2024, Supplementary Table 1). Collectively, we compiled 319 interacting structures derived by x-ray crystallography or HNMR. Compiled structures lacked the constant region of the TCRαβ complex, due to it not interacting with the pMHC. We intentionally only used human and mouse TCRαβ sequences specific to MHC class 1 and 2 molecules which encompassed a wide variety of peptides and MHC alleles (Supplementary Figure 1a-h).

### Experimental database preparation

We probed the Immune Epitope Database^24^ (IEDB; downloaded September 18, 2024) for TCR and pMHC sequences that have been experimentally confirmed to lead to an immune response but do not have existing crystal structures. In total we identified 223,691 potential positive interacting structures and 1,674 potential negative interacting structures. We filtered all sequences for human TCR structures that 1) had full length TCRαβ sequences and, 2) were human specific and, 3) matched alleles found within ground truth structures, resulting in 441 positive interacting structures and 73 negative interacting structures (Supplementary Table 2).

### Fake TCR-pMHC structural database preparation

In order to generate a library of high probability negative interacting structures, we used random TCR switching of TCR pMHC based on previous approaches to the problem^17^. 1000 randomized TCR and pMHC pairs were generated from ground truth and experimental TCR-pMHC structures. Levenshtein distance between the original set and the new TCR was calculated per pMHC molecule as previously described^25^. Pairs with Levenshtein distance less than the 90^th^ percentile of natural Levenshtein distance within a set of TCR’s specific to the same pMHC were discarded resulting in 600 falsely interacting structures (Supplementary Table 3).

### Model structural inference

Using custom scripts based on the ColabFold environment^26^, prepared sequences were batch folded on google colab servers using high-ram settings and T4 or A100 GPU’s. Alphafold2-multimer^27,28^ is an extension of the deep-learning model that uses Multiple Sequence Alignment (MSA) to improve protein folding prediction. RoseTTAfold^29^ uses MSA and physics models based on Rosetta algorithms to predict the correct full atom structure of proteins. ESMfold^30^, is an extension of the protein language model ESM to predict the correct contact sequences. All models were predicted with 5 recycles and Amber side chain relaxation. Structures were also predicted using a sequence-similarity, template-based algorithm (TCRpMHCmodels^31^) as a control to predict structures of 201 ground truth and 467 experimental MHC-I bound structures.

### Structure processing and feature extraction

After structure prediction, structures were run through a pipeline that prepared them for further analysis. Residues were normalized using PDBfixer and hydrogens were added to all heavy atoms. Where present, predicted structures were aligned to ground truth using pyMOL v3.1, otherwise aligning to the 1ao7 structure to normalize rotation and translation. Structures were then run through an in-house pipeline to extract structural, physical, and quality features from each PDB file. Detailed methodology for feature extraction is available as Supplementary Methods and python scripts for extraction are available in the project GitHub.

### Dimensionality reduction and machine learning

Features identified in predicted experimental structures were compiled into a single dataset to explore the predictability of structural features. Principle component analysis (PCA) of extracted structural features was performed in R using the stats package. Results were visualized with the factoextra package (v1.0.7.999). A classifier was trained to predict binary interaction using several classifier models (Gradient boost, Bagging, Support Vector Machine, Multilayer Perceptron, Generalized Linear Model, and Shrinkage Discriminant Analysis) from the caret R package^32^. Models were trained to make predictions on a random 70-30 train-test split with 10 cross fold validation of the train set and hyperparameter testing. Accuracy was calculated using a probability threshold of 0.5 and the Area Under the Receiver Operator Curve (AUROC) was calculated using the pROC package^33^.

### Deep learning interaction prediction

TCR-pMHC interactions were predicted using sequence and structure based inputs. For sequence based inputs, CDR3α and CDR3β sequences with their corresponding MHC alleles and epitope sequences were trained using NetTCR2.0^25^ which is a 1D Convolutional Neural Network (CNN) that predicts positive or negative interaction. To match this format, we developed a 2D CNN with matching architecture (Supplementary Figure 4, Supplementary Table 4) using the TensorFlow library. The model took as input a left-right padded contact map of dimensions 1024×1024 with the n-terminal of the epitope set at pixel 400 to normalize different sized contact maps. To combine sequence and structure information we tested different encoding methods^34^ in a different 2D CNN developed on the Pytorch library. For each cell of the contact map, the encoding for paired residues were subtracted and the Cα distance was appended to the end of the vector. A chain-pair one hot encoding was also appended to the vector and then the encoding was reconstructed into the final 2D structure of the same dimensions as the contact map and padded right to 1,024×1,024 pixels. Blosum62 encodings were selected for lowest loss so the final dimensions of each residue pair was 25 (20 dimensions for paired blosum encoding, 5 dimensions for chain encoding, 1 dimension for distance value). For all methods, we used a 70-30 train-test split trained over 100 epochs using an A100 GPU.

### Molecular dynamics (MD) and steered MD simulation

TCR-pMHC structures from ground truth, positive interacting, and fake datasets containing over-represented HLA-A2 epitopes (“ELAGIGILTV”, “SLLMWITQV”, and “SLLMWITQC”) were prepared for GROMACS^35^ molecular dynamics simulation. Based on previous work^36,37^, we ran the simulation at 100 ns with 4 fs time steps in a box environment at 300 K. Each structure was solvated in an aqueous environment with positive and negative ions. Using a Charmm27 forcefield, we used energy minimization to relax the structure and then equilibrated for temperature and pressure. During the simulation, we recorded electrostatic and RMSD changes and then calculated hydrogen bonds and energy interactions in python with the MD analysis package^38^. Dissociation energy was estimated using the gmx_MMPBSA package^39^ every 1 ns. Steered MD simulation (SMD) was performed using the VMD package^40^. Equilibrated positive and negative interacting structures were simulated for 100 ps at 2 fs timesteps. The TCR c-terminal ends were fixed at starting positions and a 5 kcal/mol spring constant was applied to 6 residue backbone Cα atoms at the bottom of the MHC (residues 6,10,25, 97,105,115) to apply an even and constant downward force at a velocity of 0.002 Å /timestep. Settings used to run all simulations can be found in the project GitHub (project/simulations/Settings/) mdp files.

### Immrep23 folding and prediction

In order to test the 2D CNN model on a dataset for which it has not learned any representation, we performed inference of the trained model on the Immrep23 test set^17^. In total, 1,022 structures were folded using AlphaFold2 and processed as above. Contact maps for 682 structures containing 9 AA epitopes (to directly compare with NetTCR) were processed as above and run on the padded 2D CNN. Extra processing was applied to the contact map to remove residues 1-24 and 204-365 from the MHCα, 1-20 and 225-272 from the TCRα, and 1-19 and 262-311 from the TCRβ due to discrepancies in sequences to the training set. We then predicted the interaction of the 682 CDR3a and peptide sequences using NetTCR^25^ (trained on the same dataset as the 2D CNN or the larger MIRA dataset) and ERGO II^41^ (both LSTM and Autoencoder).

### Statistics

Non-parametric wilcoxon ranked sum tests were used to calculate the statistical likelihood between paired comparison groups. All statistical analysis was performed in an R coding environment.

### Data and Code availability

All code is made publicly available through the project GitHub page (https://github.com/RobbenLab/TCRSIP) and requests for additional information will be made upon request.

## Results

### AlphaFold produces the highest fidelity TCR-pMHC multimeric structures

To establish a methodology for producing structural features predictive of interaction, we compared three deep-learning based models (AlphaFold2-multimer^28^, RoseTTAFold^29^, ESMfold^30^) and one sequence-homology, template-based structure prediction tool (TCRpMHCmodels^31^). We performed structural inference on 316 TCR-pMHC pairs with experimentally validated crystal structures (Fig 1a). Average inference time per structure was 12 min for AlphaFold2, 5 min for RoseTTAFold, and 1 minute for ESMFold (Fig 1b). Quality, positional, and physics features were extracted from inferred structures to compare to ground truth data (Supplementary Figure 1).

**Figure 1.**
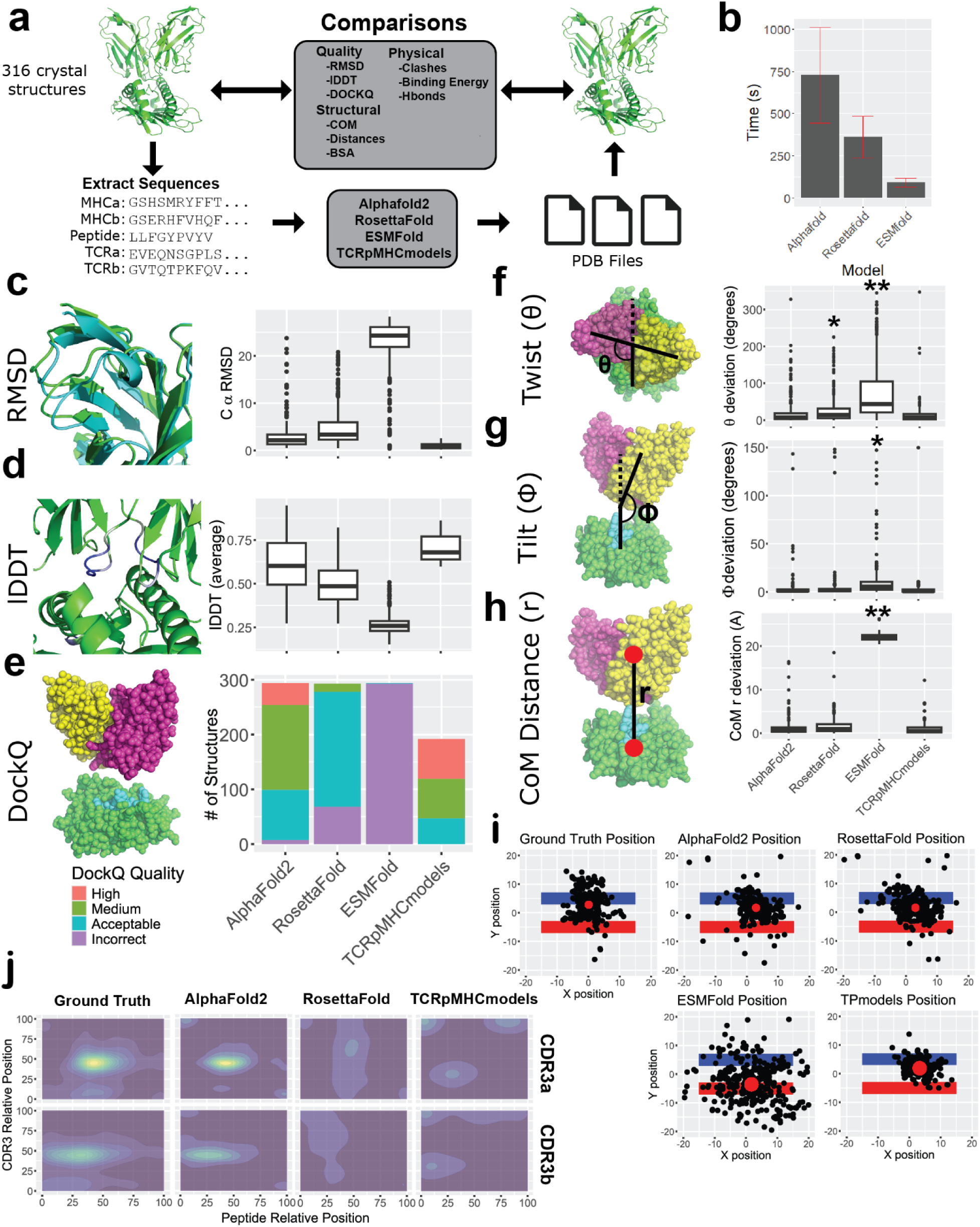
Evaluation of Quality and structural features predictive of TCR-pMHC interaction. **(a)** In total, 316 ground truth were batch input as multimeric sequences to AlphaFold2, RoseTTAFold, ESMfold and TCRpMHCmodels to predict the native complex structure. **(b)** Average time for structural inference on A100 GPU was highest for AlphaFold2. Execution time varied based on GPU use, with a T4 GPU adding on average 180 seconds per structure. RoseTTAFold and AlphaFold2 inference time is heavily influenced by server based API calls to MMSeq2 for generation of MSA, which could be reduced by pre-generated or dedicated MSA generation. Time did not vary in CPU based TCRpMHCmodels inference (∼240 seconds). **(c)** RMSD and **(d)** global lDDT calculated as average of pairwise deviation and lDDT, respectively. **(e)** DockQ classification of multimeric binding quality based on previously suggested cutoffs^35^. COM calculations were used to derive the angles **(f)** theta, **(g)** phi, and **(h)** distance at which the TCR sits over the pMHC. Reporting deviation from crystal structure measured by Wilcox signed rank test (**\***p < 0.05, **\*\***p < 0.01). **(i)** The TCR center of mass plotted from a top down view over the α1 (blue) and α2/β1 (red) arms of the MHC molecule. **(j)** Density plot of relative contact positions of the CDR3 region of the TCRα and TCRβ and peptide. Relative position is percent distance from the n-terminal side of the region on each structure.

Of the deep learning models, AlphaFold2 showed the lowest RMSD, highest lDDT, and highest proportion of “high quality” DockQ structures (Fig 1c-e, Extended Data Fig 1-e). The effect of this seemed to be a bias in AlphaFold towards the average TCR superposition, and not generalizing to more variable structures (Fig 1f-i, Extended Data 1f). The template-based algorithm underperformed AlphaFold2 in predicting peptide position, BSA, and hypervariable region positions (Fig 1j, Extended Data Fig 1c and 1g, Supplementary Fig 2). From these results it seem that AlphaFold produces structures with the most consistency while also having the flexibility to accurately position hypervariable regions.

### Physics measurements of AlphaFold produced structures fit real world structures

The strength of the T cell response has been found to be linearly correlated with the binding affinity of the TCR to the peptide MHC complex^42^. A survey of methods for predicting TCR binding free energy (ΔG) show that machine learning based approaches can predict roughly linear estimates to real world values (Fig 2a, Extended Data Fig 2a-b, Supplementary Fig 1). AlphaFold2 and TCRpMHCmodels showed the least deviation in ΔG compared to ground truth structures (Fig 2b). Surprisingly, full atom models from RoseTTAFold yielded the highest amount of high energy clashes compared to ground truth structures (Fig 2c). Hydrogen bonds (Hbonds) formed between the TCR and pMHC stabilize the immune synapse (Extended Data Fig 2c-d, Supplementary Fig 1)^43^. We found that AlphaFold produced roughly the same number of Hbonds as the ground truth, while template-based approaches could not find any between molecules (Fig 2e, Extended Data Fig 2e-g). While the number of Hbonds were constant, it appears that AlphaFold2 predicted different acceptors and donors between the CDR3 and peptide atoms (Fig 2f).

**Figure 2.**
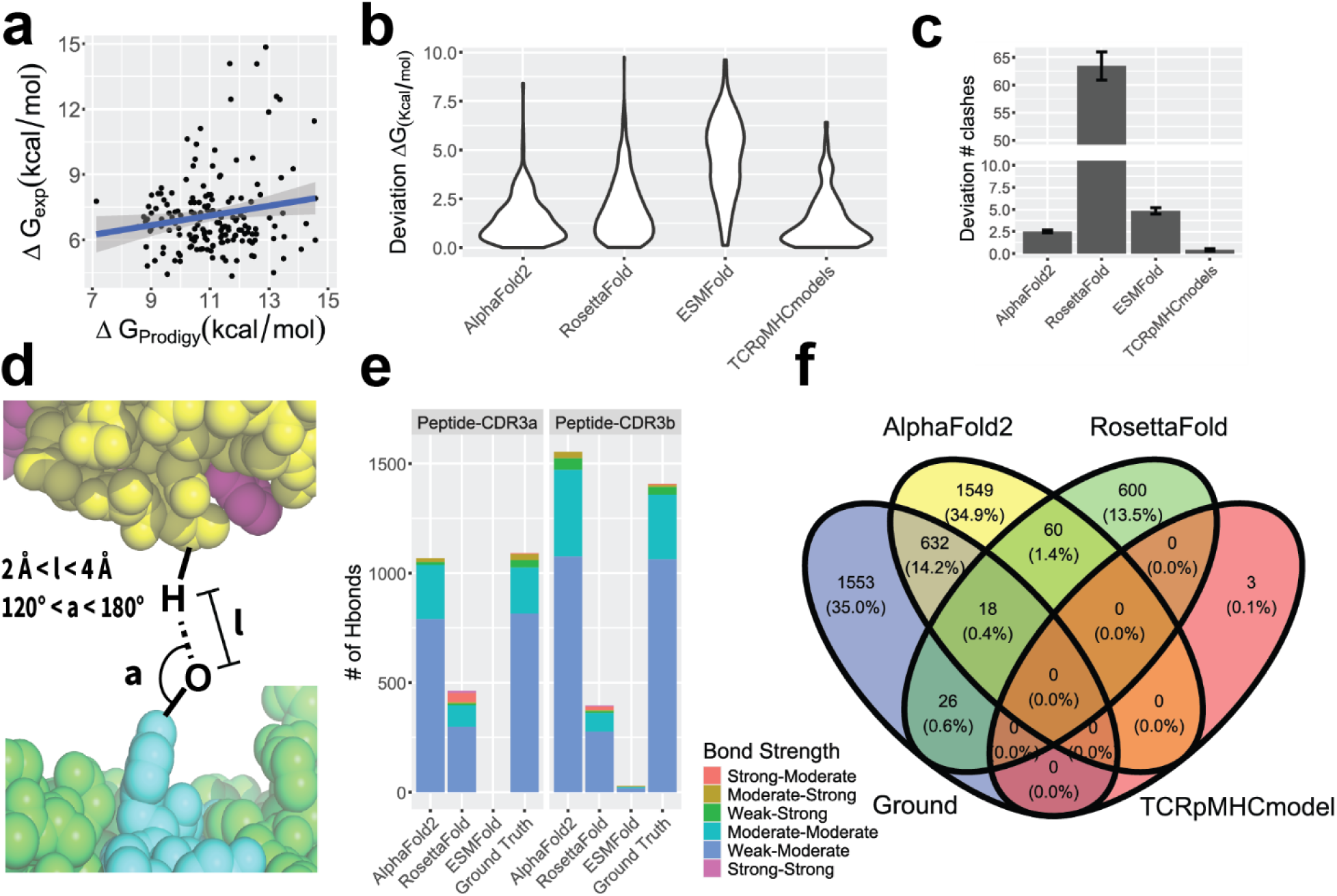
Comparison of physically based measurements between models. **(a)** Prodigy binding free energy prediction, a machine learning based algorithm, was the most consistent measure with a R^2^ value of 0.028 to real world measurements. **(b)** Deviation of prodigy measurements was calculated by subtracting prodigy scores for each ground truth structure from the value of each paired structure derived from a model. **(c)** Clashes were determined for heavy atoms in structure by overlap of atomic radii. Deviation from ground truth number of clashes was predicted per ground truth structure. **(d)** Hydrogen bonds were determined in GROMACS solvated environments and scored by bond length (l) and bond angle (a). Represented in legend as “length-angle” bond strength. **(e)** Total bond number between interacting regions was determined for the epitope interacting regions of the TCR molecule (CDR3α/β). **(f)** While similar in number, AlphaFold2 had a only about 20% similarity of exact bonds as shown via the Venn diagram.

### Predicted non-interacting structures differed from interacting structures by structural and physical features

Taken together, it is clear that quality, structure, and physics measurements are highly affected by model performance, so we chose to continue using AlphaFold2, which was the most consistent for all measures. We next sought to determine which measures can be differentiated between experimentally validated interacting and non-interacting structures. We curated a dataset of 514 experimentally validated positive and negative interacting structures from the IEDB database^44^ with no overlap to ground truth (Fig 3a). Structural inference showing little difference between classified positive and negative interactors (Fig 3b, Extended Data Fig 3a) and the existence of dual records for negative CDR3 sequences leads us to believe that “negative” sequences are ambiguous at best. We generated 600 additional non-interacting fake structures by randomly matching TCR and pMHC sequences (Fig 3c). Fake structures showed bias towards MHC-I and HLA-A alleles due to database overrepresentation but had mostly expected proportions of base MHC alleles (Fig 3c-e, Extended Data Fig 3b). Random sampling minimized Levenstein distances to real structures, reducing the possibility for false negatives (Fig 3f, Extended Data Fig 3c).

**Figure 3.**
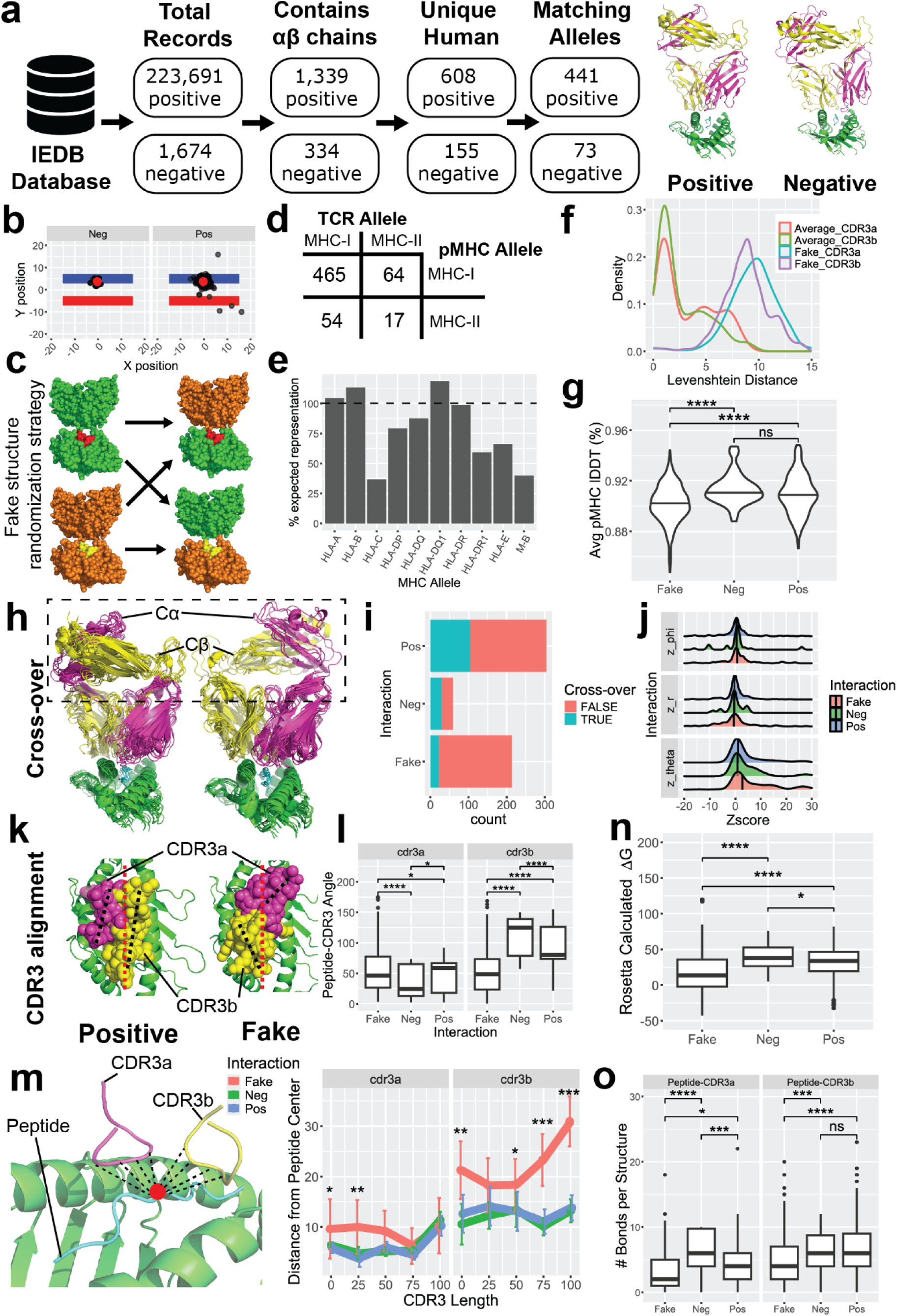
The effect of positive or negative interaction on TCR-pMHC structural features. **(a)** Dataset was generated by filtering experimentally positive and negative interacting sequences in the IEDB database for full length, human TCRab sequences that matched allelically to MHC alleles present in the ground truth dataset. An example structure for positive and negative structures is shown for the MHC HLA-A*02:01 bound to the peptide “SLLMWITQC” which is found overrepresented in the experimental dataset (14 positive interactors, 2 negative interactors). **(b)** Negative structures were found to closely match positive structures, and >80% had CDR3 sequences found elsewhere in IEDB, making their interaction ambiguous. **(c)** “Fake” structures were randomly generated by pairing orphan TCR’s to other pMHC molecules in an unbiased manner to generate example TCR-pMHC structures unlikely to interact. **(d)** Fake structures were biased towards MHC class I and HLA-A2 combinations because of the overrepresentation of these alleles in the experimental and ground truth data (∼70% of all structures).**(e)** Based on the proportion of expected alleles in the datasets, we did not find significant over-representation of most alleles compared to ground truth and experimental structures. **(f)** Comparing the Levenshtein distance of CDR3 sequences between fake and positive interactors for each pMHC against the average Levenshtein distance between all positive TCR’s for the same pMHC. **(g)** Average lDDT of the pMHC residues compared to the ground truth structure of each pMHC for negative, and positive interactors. Overlays of the positive and negative interacting structures for the pMHC HLA-A*02:01 bound to the peptide “SLLMWITQC” showing the **(h)** constant regions (Cα, Cβ) and the **(k)** CDR3. “Crossover” of the constant regions was defined by a constant region vertically over the opposite chain variable region as opposed to being over the same chain variable region. **(i)** A greater proportion of non-interacting structures showed constant region crossover. **(j)** Ridge plot showing Z-score normalized deviation from the average distance (r), twist (theta), and tilt (phi) values of structures with matching pMHC. Black vertical lines represent the population median. **(k-l)** Angle formed by the peptide linear component (red dashed line) and CDR3 linear components (black dashed lines). **(m)** Per-residue distance of the CDR3 region (n-terminal to c-terminal) to the midpoint of the peptide in each structure. **(n)** Binding energy of the TCR and pMHC calculated from Rosetta force fields for each structure. **(o)** Hydrogen bonds calculated between the peptide and CDR3 residues per structure. *p < 0.05, ** p < 0.01, *** p < 0.005, **** p << 0.005.

We find the likelihood of misfolding in inferred structures (based on clearly defined structural metrics; Extended Data Fig 3d-f), was roughly 10%, with no bias against negative interacting structures. While ground truth structures did not exist for experimentally validated sequences, we found that fake structures, in comparison to positive structures, had less than 2% difference in the lDDT of pMHC’s common to ground truth (Fig 3g). Unlike Bradley^20^, we found no difference in the pLDDT of positive and negative interactors (Extended Data Fig 3g). These results suggest that we cannot expect a difference in the quality of structural inference between positive and negative interacting structures.

Structural features did not change greatly between interacting and non-interacting predicted structures (Extended Data Fig 5h-l). An interesting observation was a bias in the majority of non-interacting structures to display a “crossover” conformation of the constant region (Fig 3h) that was less common in positive interacting structures (Fig 3i, Extended Data Fig 3m). Surprisingly, this conformation does not exist in TCR-pMHC structures found in the Protein Databank, so it is unclear why AlphaFold produced it so consistently. This “crossover” seemed to result in a tendency towards a twist (increased theta) in the superposition of the TCR (Fig 3j). Non-interacting structures also greatly affected the interaction between the TCR hypervariable regions and the pMHC (Fig 3k). The most notable effect was a change in the angle and distance of the CDR3 over the peptide (Fig 3l-m, Extended Data Fig 3n). Distance between CDR2 and CDR1 and the MHC was also increased (Supplementary Figure 3). The effect was decreased complex stability and fewer Hbonds between the peptide and CDR3β (Fig 3n-o, Supplementary Figure 3).

### Structure-based features are better predictors of immune specificity than sequence-based features

We next sought to investigate whether structural features could differentiate positive and negative interacting TCR-pMHC sequences more accurately than sequence based features. Using the features extracted from previous analysis, we found that most non-interacting fake structures segregated from interacting structures (Fig 4a, Extended Data Fig 4a-b). These features were also highly predictive in machine learning models achieving accuracies up to 94% and area under the curve (AUC) greater than 0.98 (Table 1, Fig 4b, Extended Data Fig 4c-d). Feature importance showed that energy calculations, CDR2/1 – MHCα distances, and the angle formed by the CDR3β and peptide were most predictive of interaction (Fig 4c, Extended Data Fig 4e-f).

**Figure 4.**
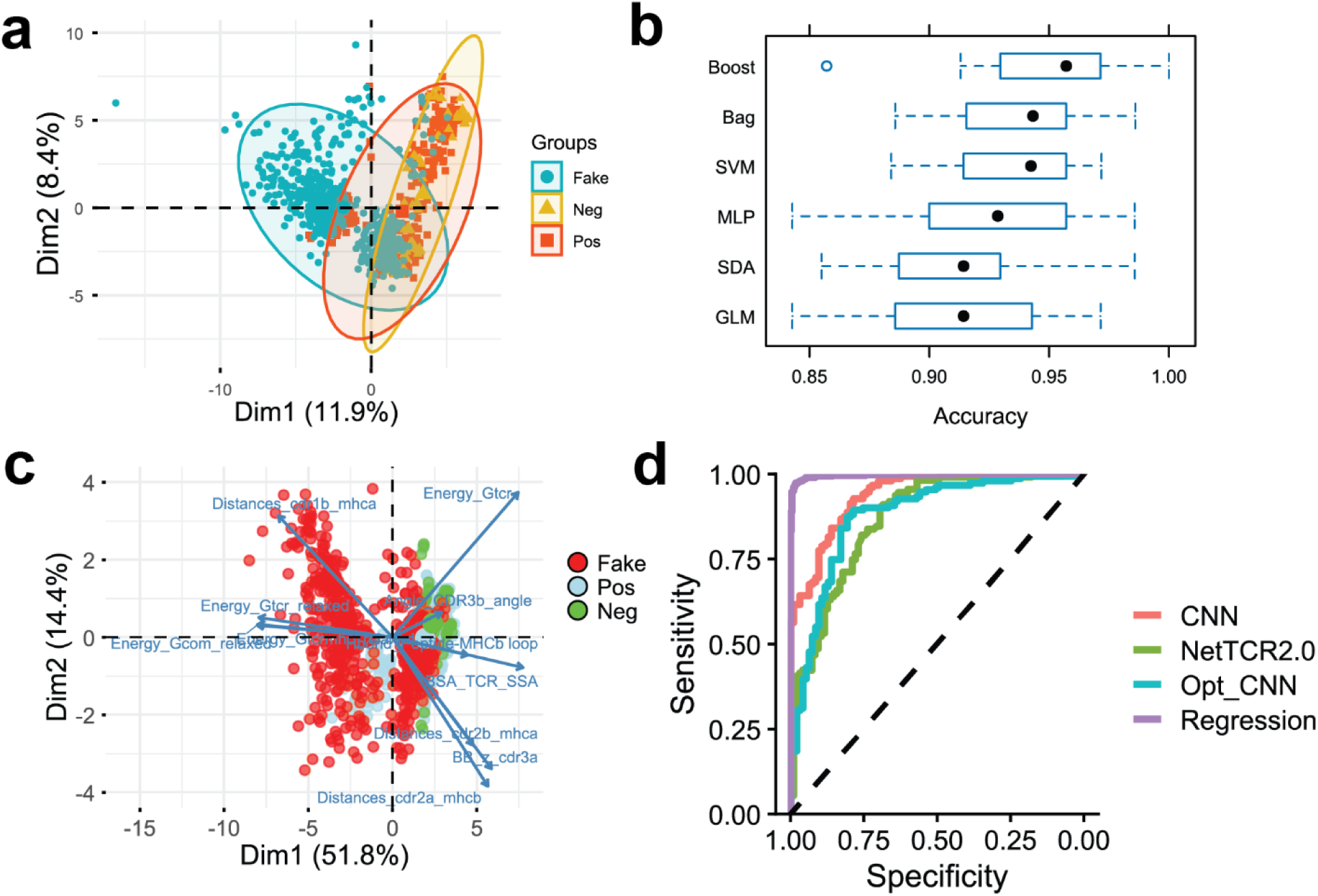
Prediction of TCR-pMHC interaction from structural features. **(a)** Principle component analysis of structural features show distance between fake structures and positive and negative structures. **(b)** Training different binary classifiers of interaction result in high discriminatory accuracy of fake from positive interacting structures. All models were trained on a 10-fold cross validation dataset. **(c)** Biplot shows the contribution of 15 most important structural features in prediction accuracy. First term, separated by underscore represents the category of type of feature for each with “Energy” features calculated from Pyrosetta forcefield scoring function. **(d)** Receiver operator curve (ROC) showing the prediction AUC of models trained on structural and sequence (NetTCR) features. “Opt_CNN” refers to the optimized architecture for a 2D CNN trained on structural features. Black dashed line showing the estimated response for a random prediction.

**Table 1.** Validation set prediction accuracies of models for sequence and spatial features.

| Model | Accuracy | AUROC | Positive Accuracy | Negative Accuracy |
| --- | --- | --- | --- | --- |
| *Extracted Features* |  |  |  |  |
| Bagging | 0.925287 | 0.983569 | 0.906667 | **0.939394** |
| SDA | 0.902299 | 0.970774 | 0.946667 | 0.868687 |
| MLP | 0.902299 | 0.970774 | 0.946667 | 0.868687 |
| GLM | 0.902299 | 0.931448 | 0.933333 | 0.878788 |
| Boost | **0.948276** | **0.988956** | **0.973333** | 0.929293 |
| *Architecture Comparison* |  |  |  |  |
| NetTCR2.0 (Sequence only) | 0.791262 | 0.869085 | 0.837838 | 0.736842 |
| 2D CNN (Structure only) | **0.860656** | **0.936766** | **0.927152** | 0.752688 |
| 2D CNN (optimized) | 0.848361 | 0.878516 | 0.874172 | **0.806452** |
| *Features comparison (combined feature CNN)* |  |  |  |  |
| Sequence CNN | 0.665768 | 0.665768 | 0.666667 | 0.666667 |
| Structure CNN | 0.590297 | 0.590297 | 0.540541 | 0.540541 |
| Sequence + Structure CNN | **0.752022** | **0.803858** | **0.797297** | **0.684564** |

We developed a 2D CNN architecture similar to the 1D CNN used in NetTCR2.0 and trained the model to predict interaction based on the input contact maps (Supplementary Fig 4). We found an increase in accuracy and AUC compared to NetTCR2.0 trained on the same dataset (Table 1, Fig 4d). The model seemed to focus on learning positionally variant features that result from padding contact maps that could prevent generalization (Supplementary Figure 4). Integrating sequence based features into unpadded contact maps was more accurate than predictions on contact maps alone (Table 1, Supplementary Figure 4).

### Simulation shows that non-interacting structures fail to stabilize the overall reaction

To observe the dynamic relationship of negative interacting structures, we generated MD simulations up to 100 ns for pMHC matched ground truth and interacting and non-interacting structures. We found that non-interacting structures had fewer stable interactions between the TCR and pMHC, with large changes in conformation of the peptide and CDR regions (Fig 5a, Supplementary Figure 5). AlphaFold seemed to predict a non-optimal starting position causing structures to search for a stable conformation (Fig 5b). However, positive interacting structures achieved this conformation quickly while fake structures continued to look for a stable conformation until the end of the simulation (Fig 5c-d). One possible cause of this was a reduced number of hydrogen bonds formed over time between interacting structures like the peptide and CDR3β (Fig 5e). We also performed steered molecular dynamics simulation (SMD) to understand how differences in structure could affect function. Perpendicular force applied to the MHC resulted in longer bond times of the interacting positive structures compared to the non-interacting fake structures (Extended Data Fig 5). TCR constant region crossover in the positive structure results in a rotational dynamic unique from uncrossed “normal” positive interacting structures and longer bond duration.

**Figure 5.**
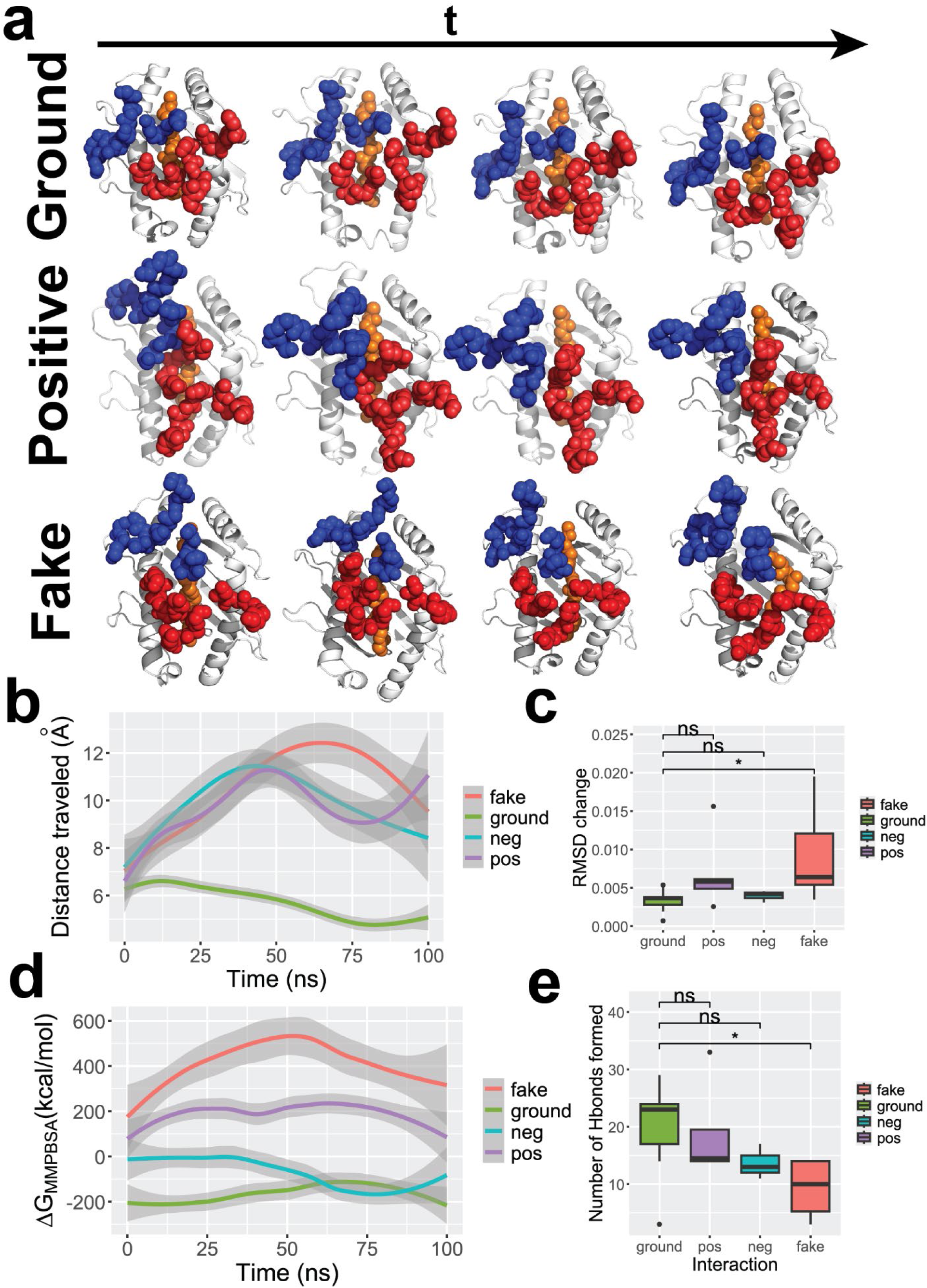
Molecular dynamics simulation of TCRpMHC structures. Structures were solvated and normalized for energy, pressure and temperature before being run for up to 100 ns with 4 fs steps. **(a)** Simulations for Ground truth (2bnq), positive interacting (pos 104), and fake non-interacting (fake 190) structures containing the peptide “SLLMWITQV” bound to HLA-A*02:01 are shown. Time steps shown at 0, 20, 40 and 60 ns for ground and positive structures and 0, 20, 40, and 70 ns for the fake structure. MHC (white) depicted as cartoon while peptide residues (orange), TCRα CDR residues (red), and TCRβ CDR residues (blue) depicted as main chain atom spheres. **(b)** Total distance traveled for interacting regions of the TCR and pMHC measured as the cumulative sum of differences between interacting regions (i.e., peptide-CDR3). **(c)** Total shift of CDR3β region shown as the cumulative difference in RMSD over all frames. **(d)** Binding energy was calculated in simulation using gmx_MMPBSA at 100 fs steps. **(e)** Number of bonds formed by individual acceptor and donors over the course of simulation is analogous to hydrogen bond extinction. Bonds were determined using minimum distance of 3.5 Å and bond angle between 120 and 180 degrees including water bridges formed between the TCR and pMHC.

### Structural features generalize predictions to unseen datasets

In order to fully assess how accurate structural features are for predicting we performed inference of interaction using an independently curated dataset from the Immrep23 competition^17^. We used AlphaFold2 to predict the structures of 682 TCRαβ sequences paired to 9 AA epitopes from the Immrep23 test dataset. The dataset encompassed 13 unique epitopes bound to MHC class I alleles with 20% experimentally validated interactors and 80% non-interacting structures. The 2D CNN model was compared to NetTCR2.0 trained on either the same dataset (Net_small) or the included MIRA dataset (Net_standard). We also compared performance to the ERGOII deep learning models^45^ using either an LSTM (Ergo_LSTM) or Autoencoder (Ergo_AE) architecture. The 2D CNN achieves higher accuracy, especially in regard to predicting negative interactions (Fig 6a-c). The prediction probabilities show relatively lower AUC, especially against the ERGO and full NetTCR models that were trained on greater than 10 times the amount of training data, likely with some overlap to the test set (Fig 6d-e).

**Figure 6.**
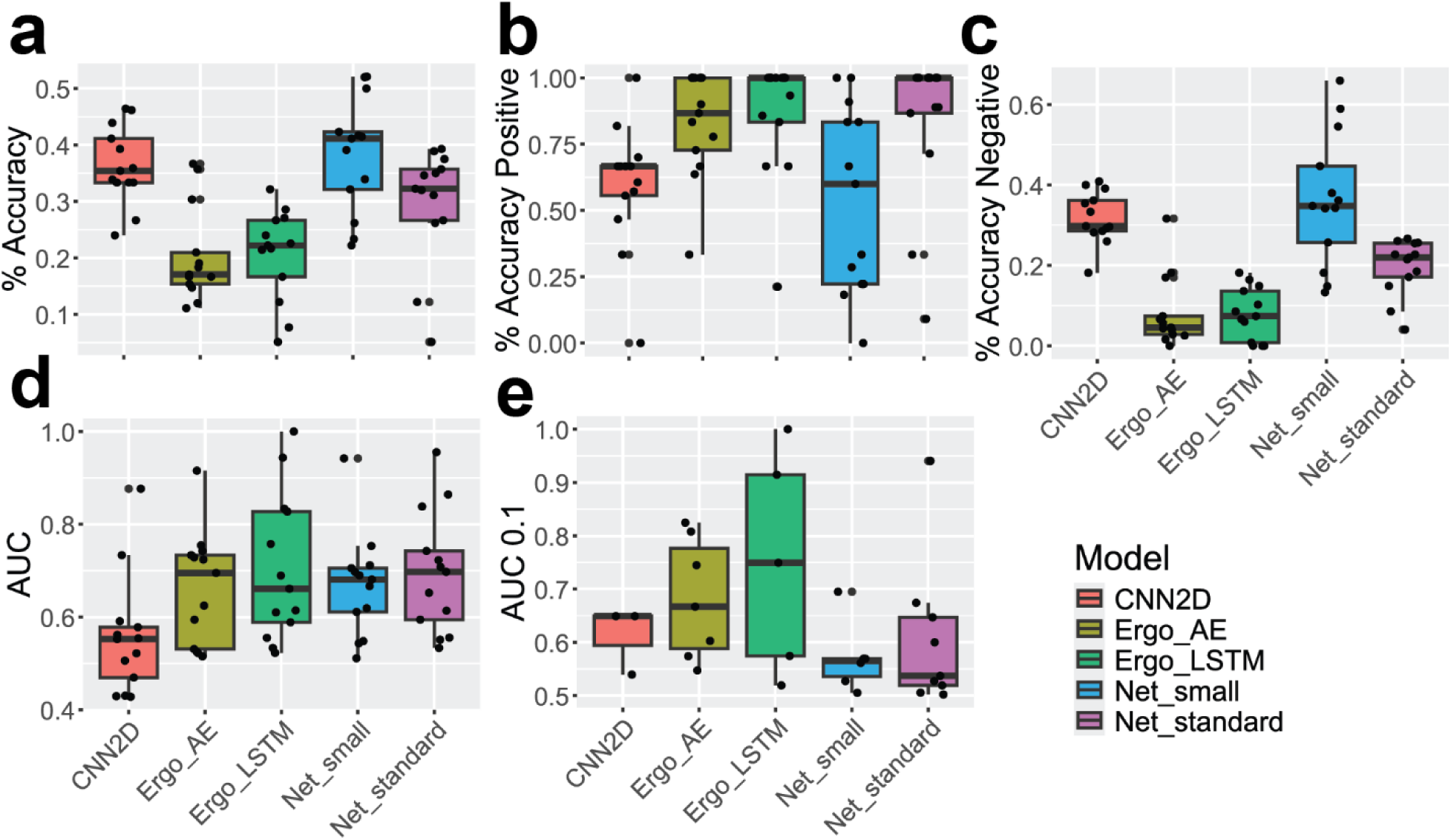
Deep learning model prediction of Immrep23 test set. In total, 682 TCR-pMHC pairs (118 experimentally positive) were used to predict binary interaction. Overall accuracy **(a)** and positive **(b)** and negative **(c)** accuracy were predicted per peptide for all structures. **(d)** AUC was calculated per peptide, and **(e)** adjusted AUC at greater than 90% specificity (AUC 0.1) was calculated for peptides for values greater than 0.5.

## Discussion

In this study we investigated the ability for structural prediction models to improve in-silico prediction of TCR-pMHC immune interactions. As a recent benchmarking study has shown^18^, the best performing sequence based models, trained on millions of data points, cannot achieve general accuracies greater than 80% on average. The compelling evidence that T cell activation is dependent upon the interaction between molecules indicates that structural data could capture information that sequence data cannot^42^. Our assumptions were validated, as structure based features demonstrated an increase in both accuracy and AUC for predicting interactions. We found that model structural inference quality itself was not a predictor of interaction, which conflicts with previous reports^20,21^. AlphaFold2 produced the highest quality structures while maintaining generalizability to unknown structures.

Extracted features that were highly predictive of interaction include hydrogen bonding, energy dynamics, and hypervariable region structure. Unfortunately, most of these features require intensive computational power to be extracted and make it difficult to use in a wide basis. Deep learning models can infer features from raw data. In comparison to NetTCR2.0^25^, a 1D CNN that takes CDR3 and peptide sequences as input, we observed higher accuracies for a 2D CNN trained on contact map data. We also observed accuracies higher in comparison to ERGOII, a more generalizable model^41^. Based on our results, we believe that structural features can be used to improve sequence based prediction algorithms and believe that such a strategy will become more common in newer approaches to specificity prediction.

We have developed a google colab webserver based implementation that incorporates AlphaFold2 structure prediction with the 2D CNN deep learning model (accessible from www.github.com/RobbenLab/TCRSIP). The advantage to this implementation is higher accuracy, prediction of the full atom structure, extraction of some structural features associated with interaction, and binary interaction prediction. The disadvantages are the time per sequence (∼20 min) which prevents the large scale screening needed for some genomic applications. Future priority in the field should be on models that could incorporate structural and physical features without expensive structural inference to make structural prediction more scaleable^46^.

Another novelty of our experimentation was the ability to observe the differences between structures that are known to interact and those that are non-interacting, as such differences may allow us to support or reject known hypotheses for TCR specificity (Fig 7). Hydrogen bonding lifetimes were increased among interacting structure simulations, as well as the time to reach an optimal energy conformation, perhaps indicating better kinetic proofreading^9^ (Fig7a) in the interacting structures. Similarly, bond strength and position differed in inferred structures, perhaps representative of a difference between catch and slip bonds^12–14^ (Fig 7b). Steered molecular dynamics simulation contributed to the idea that changes in structure resulting from a tangential or perpendicular (Fig 7c) force were responsible for activation, albeit, in a surprising and unexpected manner. Movement of the constant chain and intermembrane region was dependent on the “crossover” status of the constant chain (Fig 7d). When a constant pulling force was applied to the MHC, we observed that the bridge strands (TCRαβ residues 97∼109), would detach from the opposite side, pulling the constant regions to rotate about the z-axis. This is similar to proposed action in the force dynamics hypothesis but we only observed this mechanism in structures with “crossover” of the constant regions. If this initial conformation is so important in T cell function, it is strange that it is not present in any native TCR structures in the PDB database. We surmise that the structure itself is unstable and may require other interactors to keep it in the primed “crossed” state, which could solidify the on/off state of the TCR.

**Figure 7.**
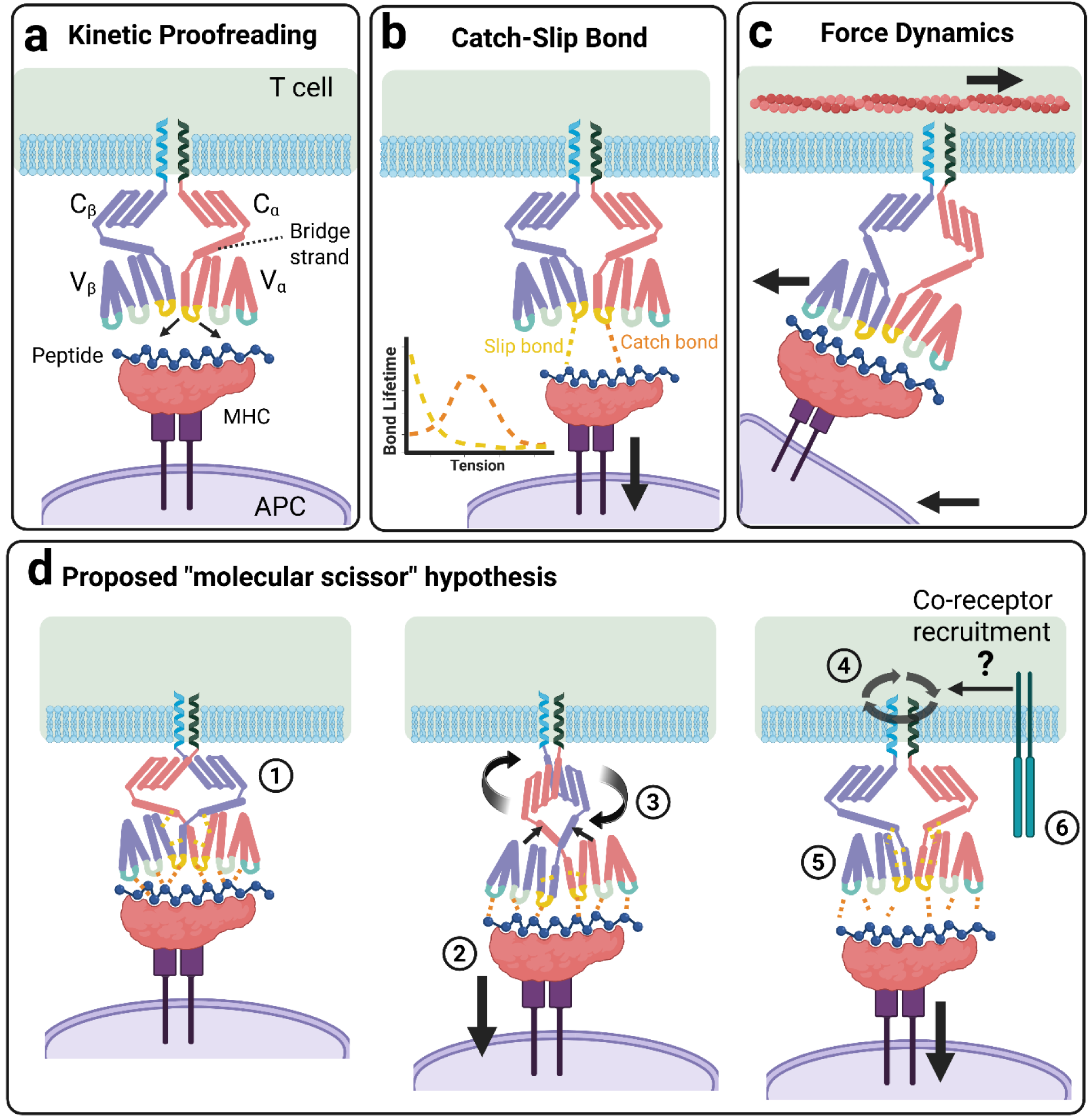
Diagram of potential molecular mechanisms that regulate TCR-pMHC specificity and activation. **(a)** Kinetic proofreading describes the multiple intermediary states that the TCR takes to reach a stable conformation. We have observed in molecular dynamics simulations that hypervariable regions (depicted as colorful loops) have increased travel in non-interacting structures. **(b)** The dynamics of catch-slip bonds, where bond lifetime increases as tension forces increase (catch bond), have been used to describe TCR specificity^14^. There is little agreement as to what bonds constitute catch or slip bonds. **(c)** The force dynamic hypothesis is based on observations that the TCR acts as a mechano-sensor, signaling downstream after a perpendicular shear force is applied to the variable region by the APC^11^. **(d)** The mechanism of TCR specificity supported by our evidence requires some aspect of each of these hypotheses. In the scissor hypothesis, the TCR structure begins naïve with a crossed constant region, secured by bridge strand interaction with the opposite chain **(1)**. Downward force **(2)** from the MHC being pulled by the APC results in the weak interchain bonds of the bridge strand breaking **(3)**, straightening the strands and forcing the constant region to rotate parallel to the TCR membrane **(4)**. The bridge strands may stabilize by bonding in the “uncrossed” conformation before or after the pMHC dissociates from the TCR **(5)**. TCR activation may be induced by the rotational movement of the intermembrane tail itself or by allowing recruitment of stabilizing co-receptors **(6)**.

## Acknowledgements

The author thanks Diwakar Shukla and Deborah Leckband for providing advice and insights regarding molecular dynamics and biophysics that improved this manuscript. The author acknowledges the USDA Hatch program for supporting the project with discretionary funds. The author also thanks the University of Illinois, Urbana-Champaign Institutional Research Board (Award #RB25159) for providing additional funds crucial for supporting this work.

## Author contributions

M.R. conceived the initial idea for the study, performed all experiments, put together figures and composed the manuscript.

---

**Supplementary Figure 1.**
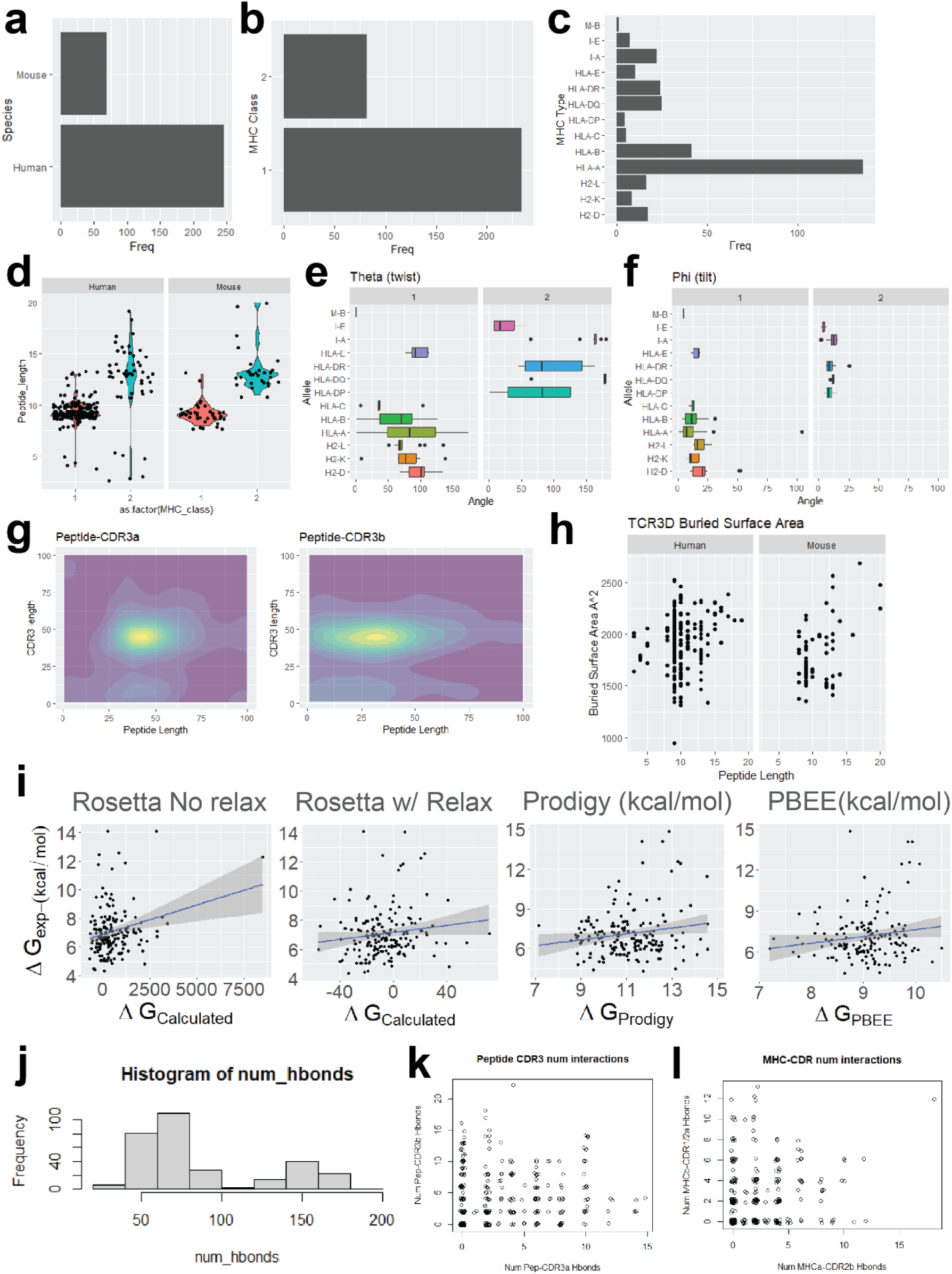
Ground truth TCR-pMHC structure characteristics were analyzed for properties that could be used as metrics of structure prediction quality and interaction features. **(a-c)** Structures were predominantly from humans and MHC-I interacting TCR structures. There was over-representation of HLA-A, specifically HLA-A2 bound TCR’s due to the overabundance of previous research of HLA-A2 bound antigens. **(d)** Peptide lengths in both human and mouse were on average 9 aa and 13 aa for MHCI- and MHCII-bound structures, respectively. **(e-f)** COM calculated values for the TCR twist (theta) and tilt (phi) over where it sits on the binding pocket of the pMHC molecule. **(g)** Structures had predominantly a single peak at which the CDR3α/β potentially interacted with the peptide which was focused in the middle of the CDR3 region based on a density map of all peaks. **(h)** Buried surface area (BSA) showed a linear relationship with peptide length, showing even coverage by interacting TCRs. **(i)** Real world calculated binding energies (derived from K_d_ measurements) were compared to multiple methods of calculation. Rosetta scoring function was used either with or without full atom relaxation prior to measurement. Prodigy and PBEE were both run using structures as input. Native structures show consistent hydrogen bonding **(j)** structure wide, **(k)** between peptide-CDR3, and **(l)** MHC arm-CDR2/1 residues. Less than 30 structures did not have either a CDR3α or CDR3β hydrogen bond with a median of 8 bonds per structure.

**Extended Data Figure 1.**
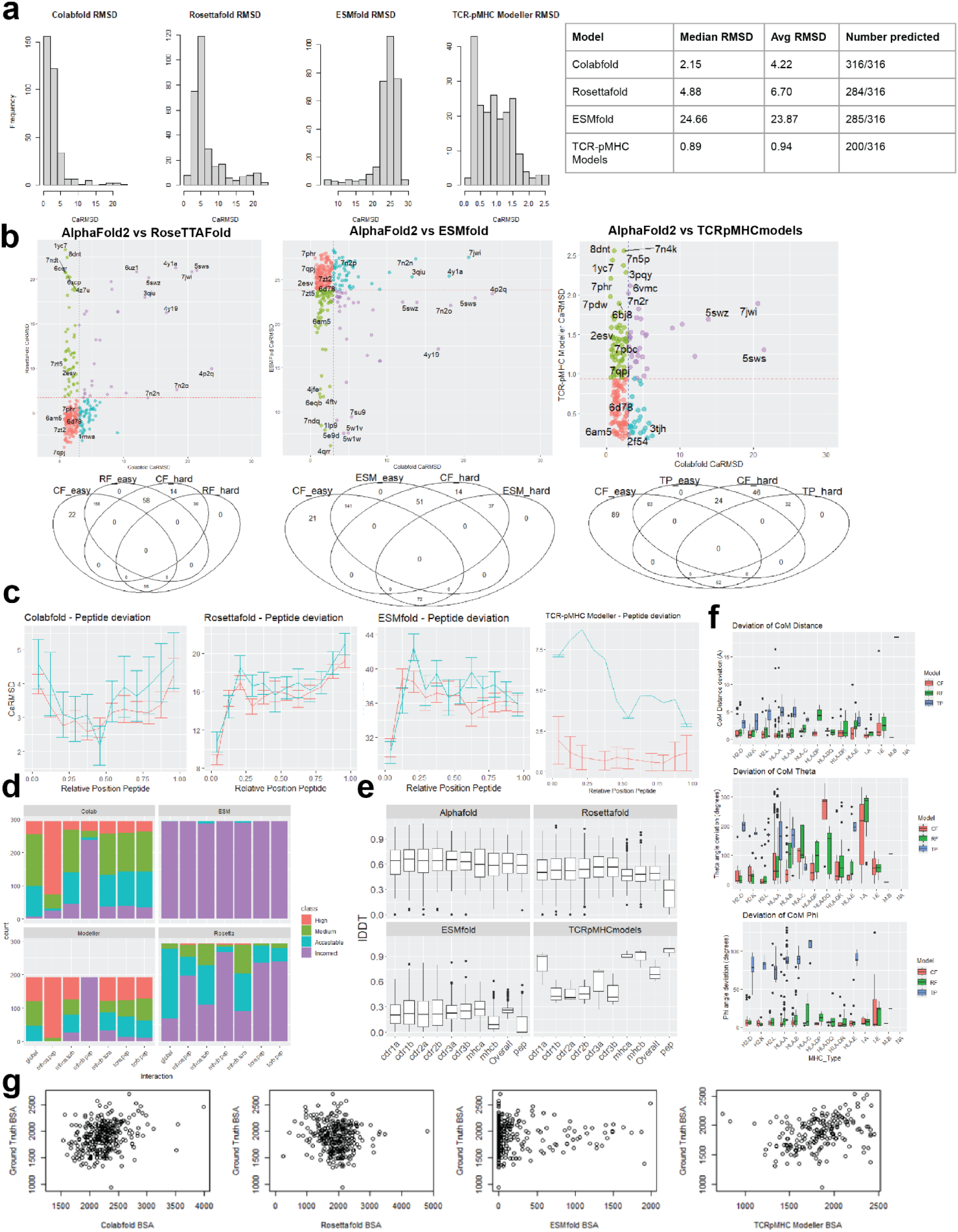
Comparisons of the structure prediction models on TCR-pMHC structural inference. AlphaFold2 and ColabFold used interchangeably in figures and abbreviated; ColabFold (CF), RoseTTAFold (RF), ESMfold (ESM), and TCRpMHCmodels (TP). **(a)** Measured global CαRMSD of different models. **(b)** Comparison of per structure RMSD performance in different models showed some universally hard to fold structures (i.e., 5swz, 7jwi, 4y1a). Difficulty of prediction defined as global RMSD over 80^th^ percentile in each model (20^th^ percentile in ESMfold due to overall poor quality). **(c)** Positional dependence of peptide quality shows lower accuracy to ends of peptide position. **(d)** Local DockQ scores for chain interactions within predicted structures. **(e)** Local lDDT prediction in each structure. **(f)** Effect of HLA type on COM scores in each model. **(g)** Linear relationship of ground truth and predicted BSA per structure.

**Supplementary Figure 2.**
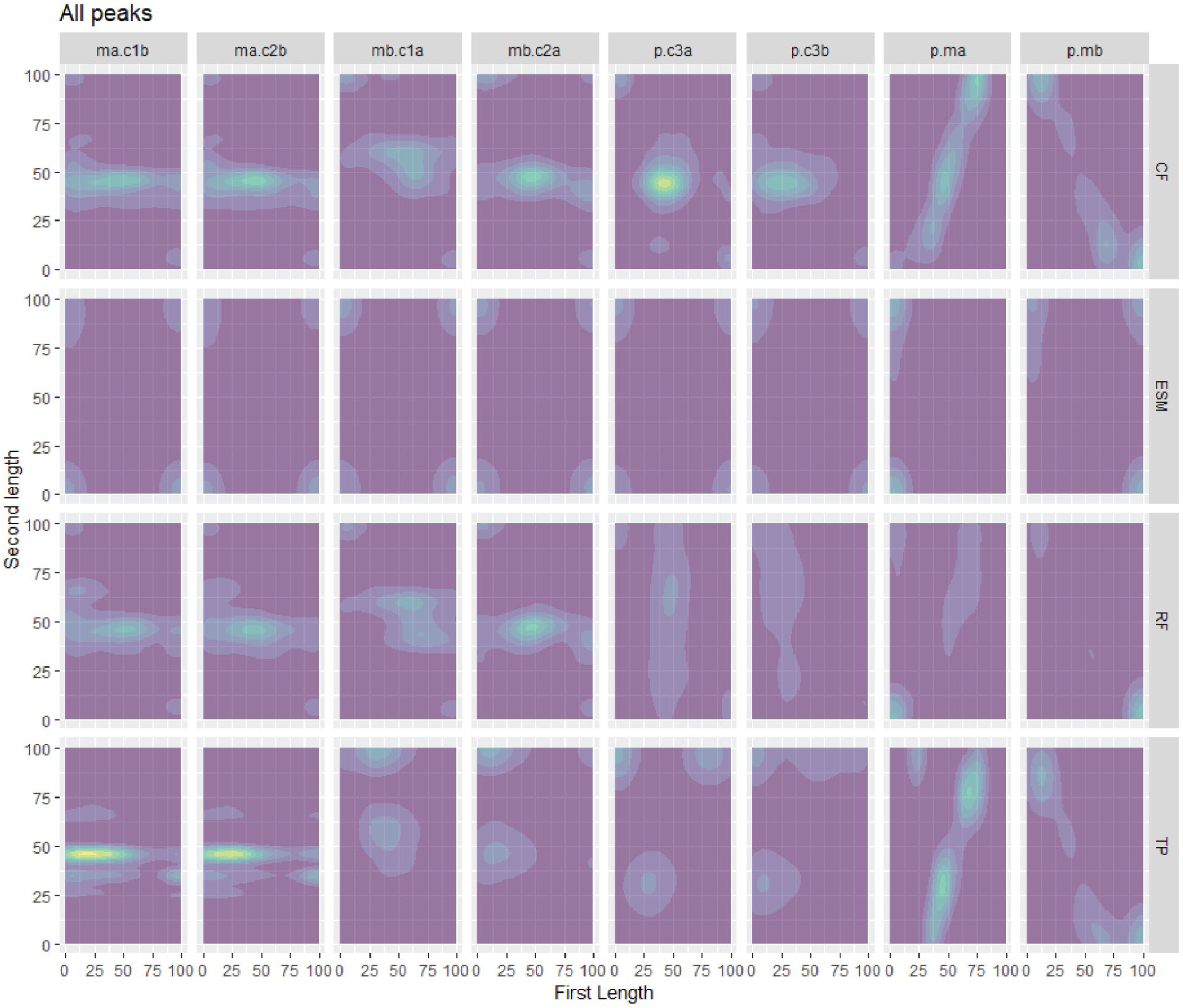
Density map of peaks for each model by region.

**Extended Data Figure 2.**
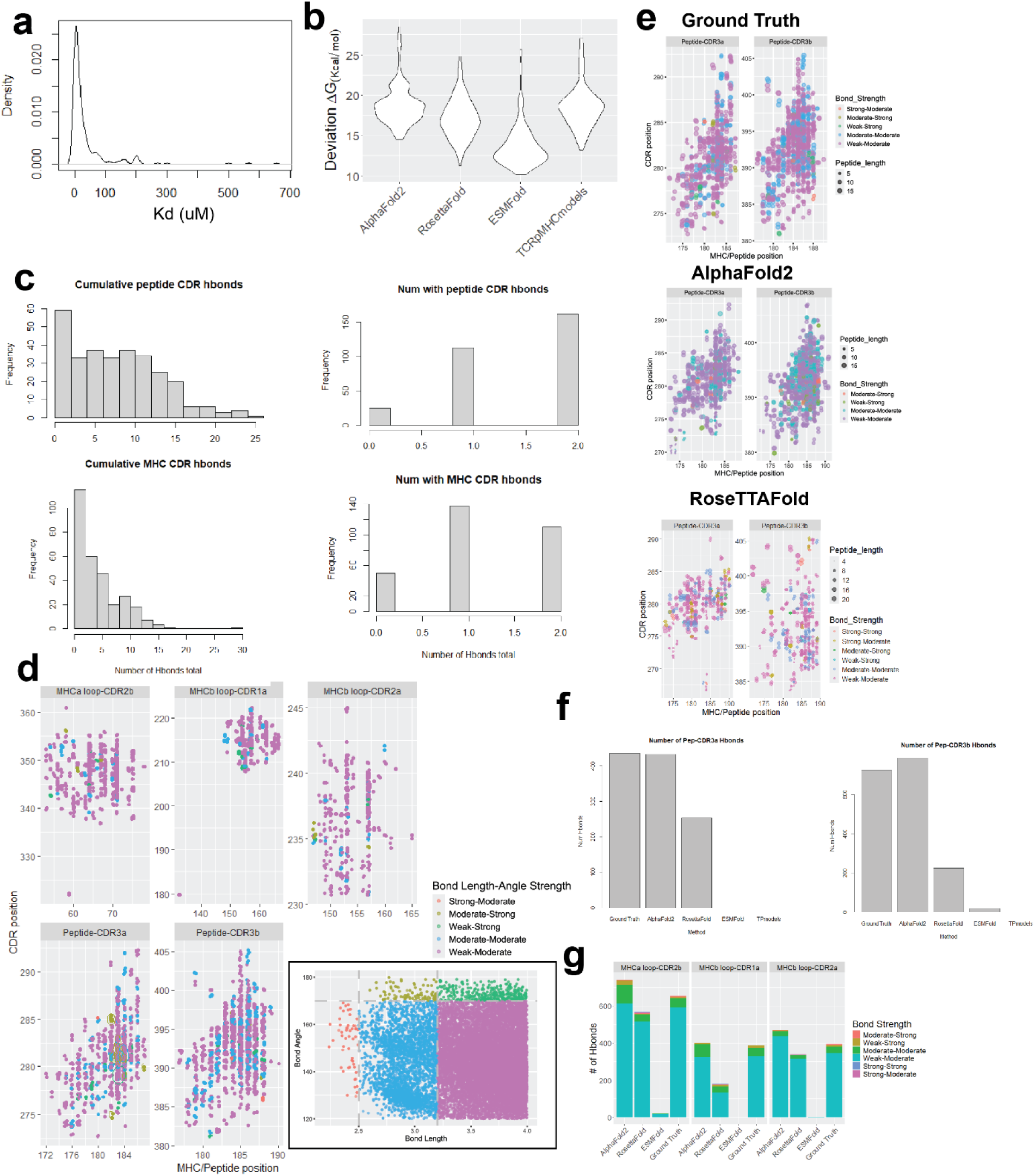
Comparative analysis of physically based measurements between models. **(a)** Density plot of experimentally validated dissociation constants (K_d_) of 316 ground truth structures. **(b)** Deviation of prodigy calculated from experimentally measured ΔG converted from K_d_ using equation (ΔG = RT ln(K_d_)). **(c)** Calculation of Hbonds in each structure between pre-defined interacting regions peptide-CDR3α/β (top) and MHCα/β loop – CDR1/2 (bottom). Total bonds found per structure (left) and structures with 0,1, or >2 bonds per structure (right). **(d)** Position of peptide/MHC residue (x axis) plotted against position of CDR3/2/1 position (y axis) by interacting region. Color representing the hydrogen bond strength by length/angle (cutout). **(e)** Comparison of peptide-CDR3αβ hydrogen bond position between 3 different models. **(f)** Number of Hbonds found in each model in the Peptide and CDR3α (left) or CDR3β (right) interaction. Number of Hbonds found in each model in MHC and CDR2/1 interaction.

**Extended Data Figure 3.**
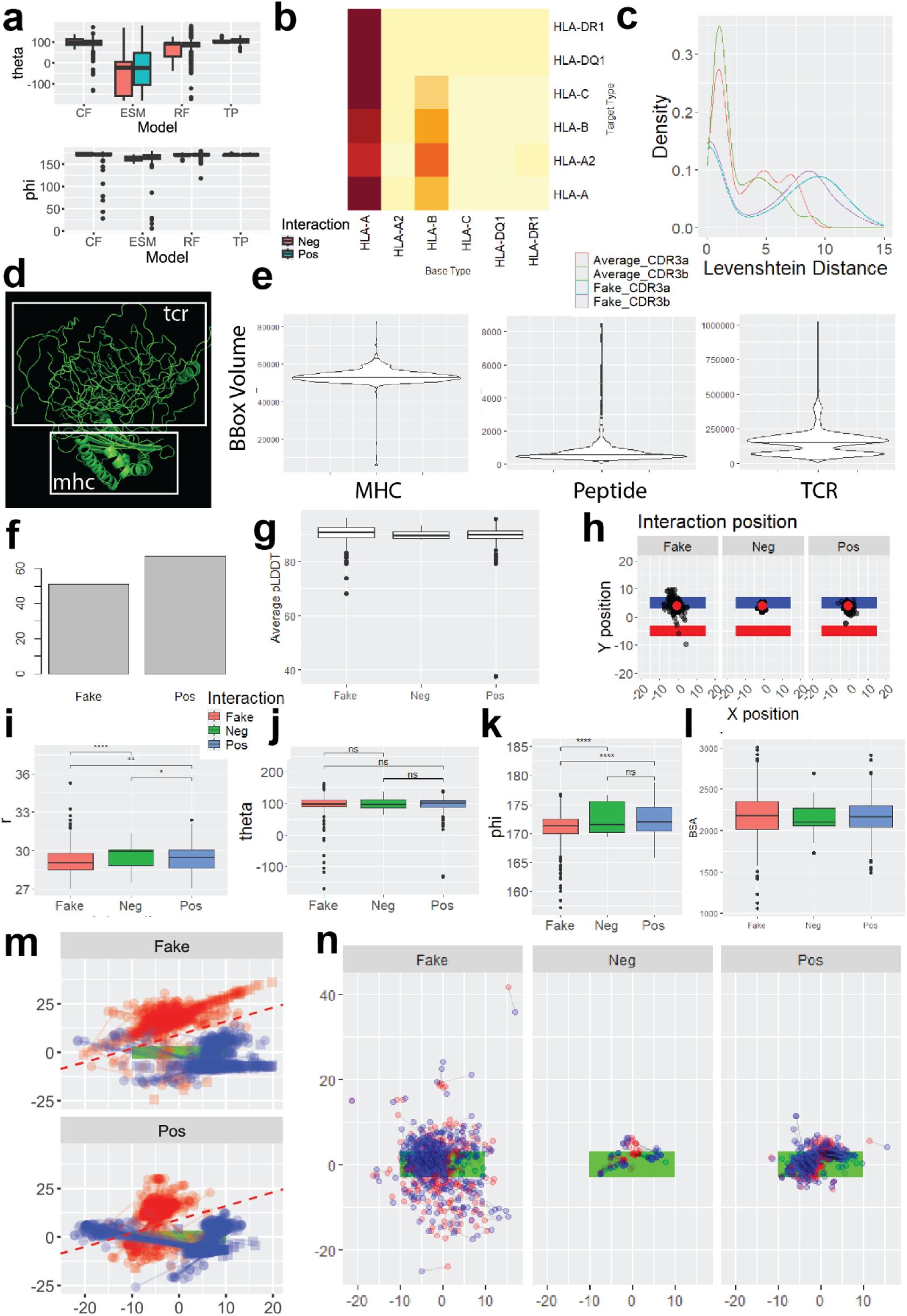
Characteristics of experimental structures. **(a)** Theta and phi angles measured of structures folded using different models (CF = AlphaFold2, ESM = ESMfold, RF = RoseTTAFold, TP = TCRpMHCmodels). **(b)** Heatmap showing the proportional mix of randomly pairing TCR and pMHC by the actual pMHC allele (base) and the allele the new TCR was previously bound to (target). **(c)** Density plot showing the Levenshtein distance for fake structures equally sampled from TCRpMHC datasets. Five structures per MHC allele TCRpMHC combination were generated and distance was determined from new CDR3 sequence to all sequences common to base allele. **(d)** Representation of misfolded protein, characterized by lack of secondary structure and poor orientation of interacting chains. **(e)** Outliers of bounding boxes of each interacting molecule were used to determine misfolded proteins with a cutoff at the 90^th^ percentile. Misfolded proteins were determined from a combination of measurements including overall volume of structure, orientation of regions, the proportion of residues within an expected region, with the overall number **(f)** of misfolded structures reported. No negative interacting structures were reported misfolded. **(g)** plDDT measurements reported by the AlphaFold2 model per structure was calculated as the average over all residues. **(h)** The center of mass of all structures in the x and y axis (top down) shows superposition of the TCR (black dot) over the MHCα1 arms (blue rectangle) and MHCα1/β2 arms (red rectangle). The average x and y position represented with a red dot. **(i-k)** The distance (r), twist (theta) and tilt (phi) of the TCR positioned over the pMHC molecule calculated by center of mass. **(l)** Buried surface area of interacting and non-interacting structures. **(m)** Plot showing the x and y position of the center of mass of the variable (square point) and constant (circle point) region of each TCR over the average peptide position (green rectangle). TCRα (red) and TCRβ (blue) chain variable and constant regions are connected by a single line. Diagonal grey line represents the cutoff used to determine “crossover.” **(n)** Plot showing the x and y position of the CDR3α (red dot) and CDR3β (blue dot) centers of mass positioned over the peptide (green rectangle). CDR3 centers of mass from the same TCR are connected by a black line.

**Supplementary Figure 3.**
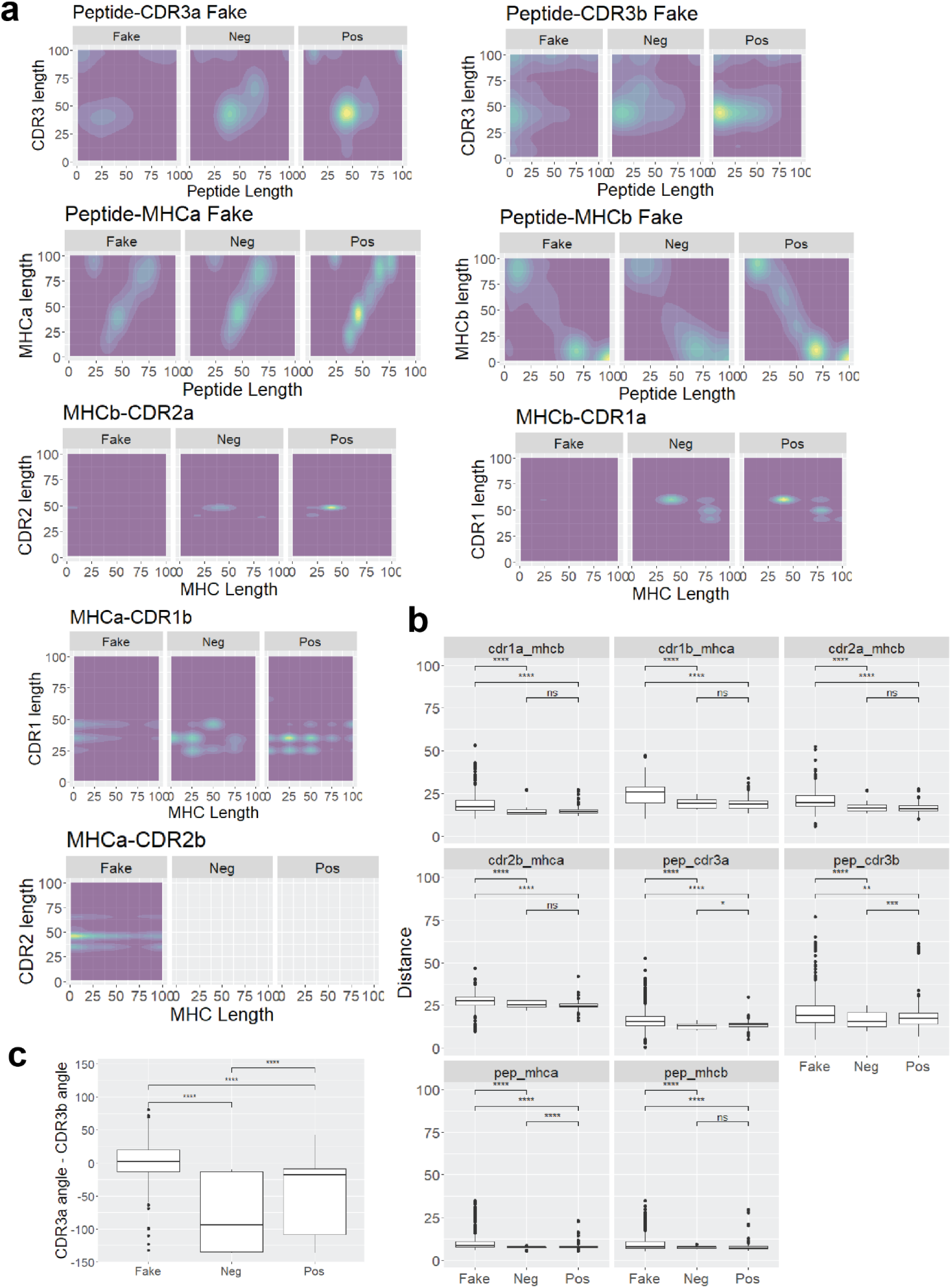
Characteristics of interacting regions of the TCR and pMHC. **(a)** 2D density plots showing the peak interaction locations of the two regions on the x and y axis. Values 0-100 represent the normalized distance (n-terminal to c-terminal) of each region. Green colors represent areas where more structures were found to have interacting residues as defined by the closest inter-residue distance at least closer than 10 Å. **(b)** Distance between the center of mass of each interacting region. **(c)** The difference between the CDR3α and the CDR3β angle shows that both regions are perturbed in the prediction of a non-interacting structure.

**Extended Data Figure 4.**
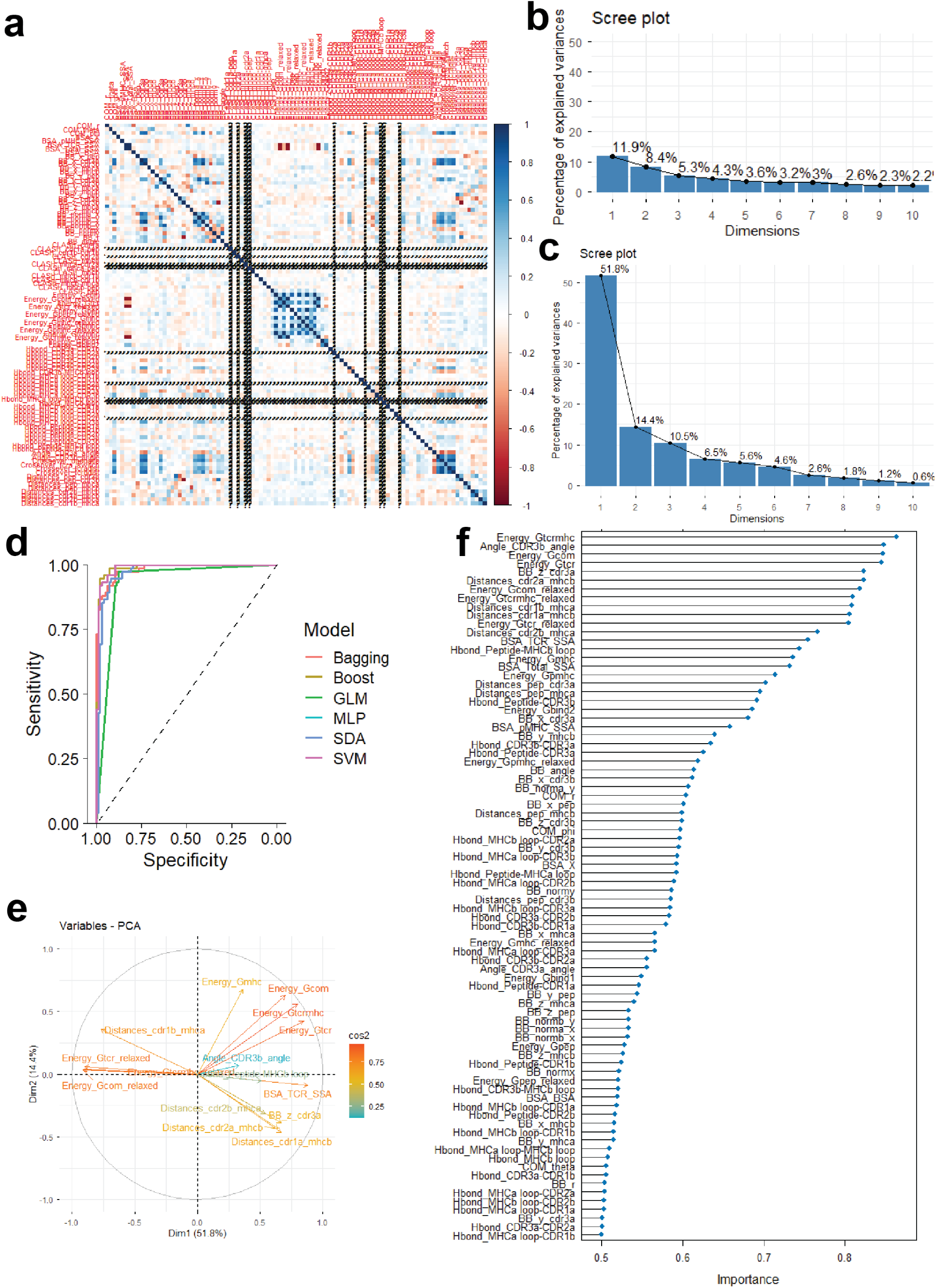
Data exploration of extracted features from predicted interacting and non-interacting structures. **(a)** Heatmap showing the correlation between extracted quantitative features. Category separated by underscore proceeding every calculated variable to differentiate extraction methods. **(b)** Variance explained in principal components between all features. **(c)** Variance increases when only considering high variance features with the highest coefficients in a logistic regression. A total of 6 machine learning models trained on 80% of applicable training data (121 structures discarded due to the presence of NA). **(d)** Receiver Operator Curves (ROC) showing the precision of model predictions in 20% validation dataset. **(e)** Biplot for variables and their effect on first two principal components with squared cosine representing the magnitude of the effect on principal components. **(f)** Feature importance map for support vector machine model.

**Supplementary Figure 4.**
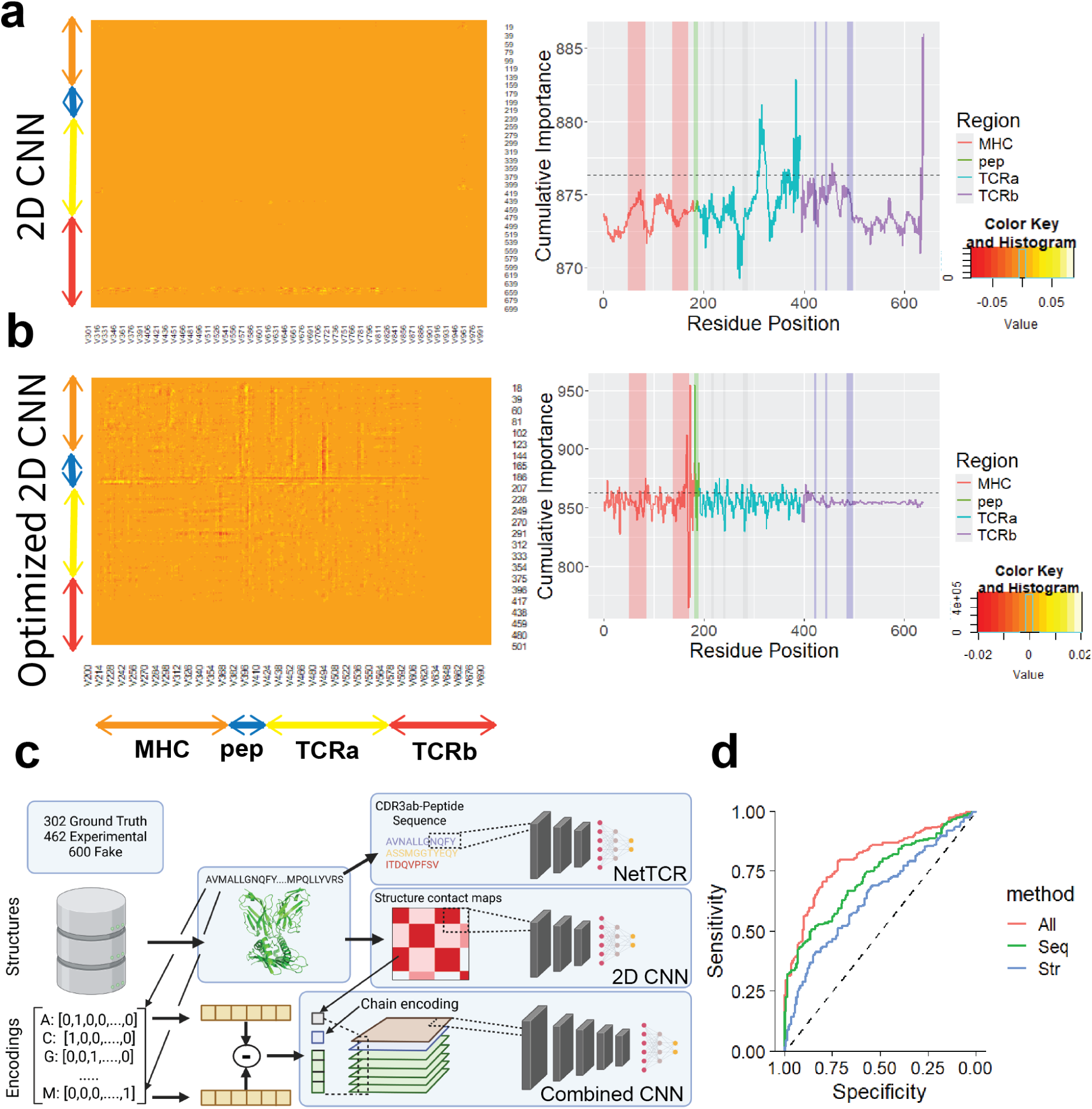
Deep learning models predict interactions between structures using contact maps. Summed feature importance of test structures is reported based on approximate chain position for **(a)** 2D CNN and **(b)** optimized 2D CNN. Cumulative feature importance at each residue position is reported on the y axis with the horizontal line plotted at the 90^th^ percentile of importance scores. Interacting regions in MHC, peptide, and TCRαβ (CDR1-3) are highlighted by vertical lines. **(c)** General strategy for developing deep learning models is reported. Sequence based features were derived from peptides for NetTCR2.0 but for the combined sequence we used Blosum64 encodings for each amino acid, using the difference between vectors for each pair. Chain encodings were one hot encoded for each chain pair and the physical distance between residue Cα’s was appended to the end of the resultant vector. **(d)** ROC of the combined CNN by which features were included. Black dashed line representing the 50% AUC.

**Supplementary Figure 5.**
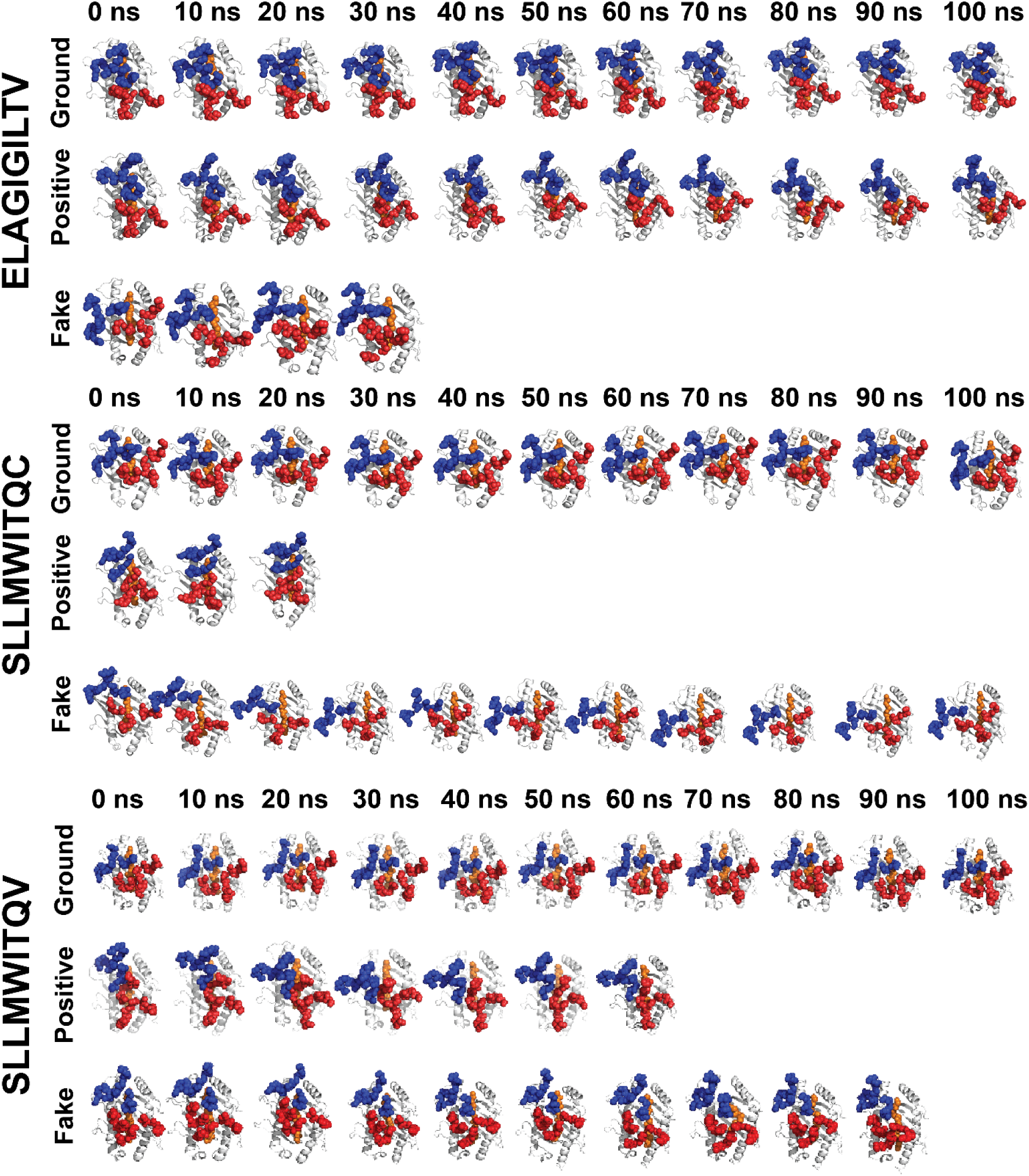
Examples of simulations run for three peptides that included ground truth, positive and fake structures. Simulations ran up to 100 ns with early stopping for LINCS errors > 1000. PDB structures for ELAGIGILTV (5nht, pos170, fake149), SLLMWITQV (2bnq, pos104, fake190) and SLLMWITQC (2bnr, pos47, fake93) were first solvated with water using a TIP3 model and NaCl ions before equilibration. All simulations were run using an amber99 forcefield in GROMACS software.

**Extended Data Figure 5.**
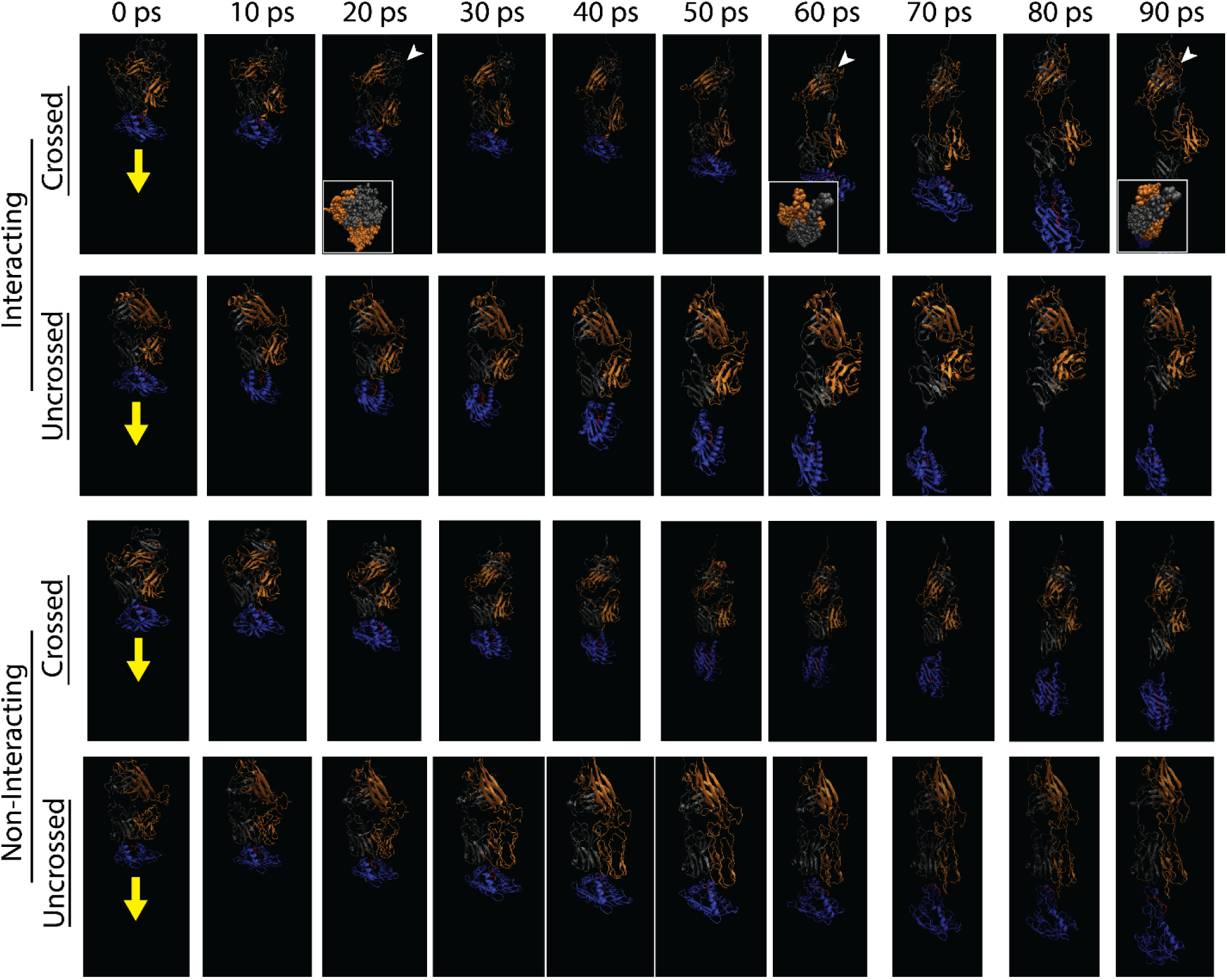
Steered molecule dynamics simulation of TCR-pMHC structures. Interacting (Pos 24136 and pos 67) and non-interacting (fake 123 and fake 10) structures were equilibrated at 300 K in a solvated environment before simulating dissociation. A downward force (yellow arrow) was applied to the MHC molecule while the C-terminal residues of the TCR were fixed into place and the simulation was run for a maximum of 100 ps. Representatives of crossed (pos 67 and fake 123) constant region structures were simulated to compare to uncrossed (pos 24136 and fake 10) structures. The constant region of pos 67 (white arrowhead) undergoes rotational motion as depicted in a top down view of the globular protein in the cutout. Structures colored by chain representation.

## References

1. Shah, K., Al-Haidari, A., Sun, J. & Kazi, J. U. T cell receptor (TCR) signaling in health and disease. Signal Transduct. Target. Ther. 6, 412 (2021).

2. Rudolph, M. G. & Wilson, I. A. The specificity of TCR/pMHC interaction. Curr. Opin. Immunol. 14, 52–65 (2002).

3. Rudolph, M. G., Stanfield, R. L. & Wilson, I. A. HOW TCRS BIND MHCS, PEPTIDES, AND CORECEPTORS. Annu. Rev. Immunol. 24, 419–466 (2006).

4. Govern, C. C., Paczosa, M. K., Chakraborty, A. K. & Huseby, E. S. Fast on-rates allow short dwell time ligands to activate T cells. Proc. Natl. Acad. Sci. U. S. A. 107, 8724–8729 (2010).

5. Shevyrev, D. V., Tereshchenko, V. P. & Sennikov, S. V. The Enigmatic Nature of the TCR-pMHC Interaction: Implications for CAR-T and TCR-T Engineering. Int. J. Mol. Sci. 23, 14728 (2022).

6. Kaitao, L., William, R., Zhou, Y. & Cheng, Z. Single-molecule investigations of T-cell activation. Curr. Opin. Biomed. Eng. 12, 102–110 (2019).

7. Szeto, C., Lobos, C. A., Nguyen, A. T. & Gras, S. TCR Recognition of Peptide–MHC-I: Rule Makers and Breakers. Int. J. Mol. Sci. 22, 68 (2020).

8. Singh, N. K. et al. Emerging concepts in T cell receptor specificity: rationalizing and (maybe) predicting outcomes. J. Immunol. Baltim. Md 1950 199, 2203–2213 (2017).

9. Gálvez, J., Gálvez, J. J. & García-Peñarrubia, P. Is TCR/pMHC Affinity a Good Estimate of the T-cell Response? An Answer Based on Predictions From 12 Phenotypic Models. Front. Immunol. 10, (2019).

10. Schodin, B. A., Tsomides, T. J. & Kranz, D. M. Correlation Between the Number of T Cell Receptors Required for T Cell Activation and TCR–Ligand Affinity. Immunity 5, 137–146 (1996).

11. Kim, S. T. et al. The αβ T Cell Receptor Is an Anisotropic Mechanosensor*. J. Biol. Chem. 284, 31028–31037 (2009).

12. Sibener, L. V. et al. Isolation of a Structural Mechanism for Uncoupling T Cell Receptor Signaling from Peptide-MHC Binding. Cell 174, 672–687.e27 (2018).

13. Hong, J. et al. Force-Regulated In Situ TCR–Peptide-Bound MHC Class II Kinetics Determine Functions of CD4+ T Cells. J. Immunol. 195, 3557–3564 (2015).

14. Faust, M. A., Rasé, V. J., Lamb, T. J. & Evavold, B. D. What’s the Catch? – The Significance of Catch Bonds in T cell Activation. J. Immunol. Baltim. Md 1950 211, 333–342 (2023).

15. Qi, F. et al. A roadmap for T cell receptor-peptide-bound major histocompatibility complex binding prediction by machine learning: glimpse and foresight. Brief. Bioinform. 26, bbaf327 (2025).

16. Hudson, D., Lubbock, A., Basham, M. & Koohy, H. A comparison of clustering models for inference of T cell receptor antigen specificity. ImmunoInformatics 13, 100033 (2024).

17. Nielsen, M. et al. Lessons learned from the IMMREP23 TCR-epitope prediction challenge. ImmunoInformatics 16, 100045 (2024).

18. Drost, F. et al. Benchmarking of T cell receptor-epitope predictors with ePytope-TCR. Cell Genomics 5, (2025).

19. Jumper, J. et al. Highly accurate protein structure prediction with AlphaFold. Nature 596, 583–589 (2021).

20. Bradley, P. Structure-based prediction of T cell receptor:peptide-MHC interactions. eLife 12, e82813 (2023).

21. Yin, R. et al. TCRmodel2: high-resolution modeling of T cell receptor recognition using deep learning. Nucleic Acids Res. 51, W569–W576 (2023).

22. Deleuran, S. N. & Nielsen, M. NetTCR-struc, a structure driven approach for prediction of TCR-pMHC interactions. Front. Immunol. 16, (2025).

23. Lin, V. et al. TCR3d 2.0: expanding the T cell receptor structure database with new structures, tools and interactions. Nucleic Acids Res. 53, D604–D608 (2025).

24. Vita, R. et al. The Immune Epitope Database (IEDB): 2024 update. Nucleic Acids Res. 53, D436–D443 (2025).

25. Montemurro, A. et al. NetTCR-2.0 enables accurate prediction of TCR-peptide binding by using paired TCRα and β sequence data. *Commun*. Biol. 4, 1–13 (2021).

26. Mirdita, M. et al. ColabFold: making protein folding accessible to all. Nat. Methods 19, 679–682 (2022).

27. Evans, R. et al. Protein complex prediction with AlphaFold-Multimer. 2021.10.04.463034 Preprint at 10.1101/2021.10.04.463034 (2022).

28. Jumper, J. et al. Highly accurate protein structure prediction with AlphaFold. Nature 596, 583–589 (2021).

29. Baek, M. et al. Accurate prediction of protein structures and interactions using a three-track neural network. Science 373, 871–876 (2021).

30. Lin, Z. et al. Evolutionary-scale prediction of atomic-level protein structure with a language model. Science 379, 1123–1130 (2023).

31. Jensen, K. K. et al. TCRpMHCmodels: Structural modelling of TCR-pMHC class I complexes. Sci. Rep. 9, 14530 (2019).

32. Kuhn, M. Building Predictive Models in R Using the caret Package. J. Stat. Softw. 28, 1–26 (2008).

33. Robin, X. et al. pROC: an open-source package for R and S+ to analyze and compare ROC curves. BMC Bioinformatics 12, 77 (2011).

34. Jing, X., Dong, Q., Hong, D. & Lu, R. Amino Acid Encoding Methods for Protein Sequences: A Comprehensive Review and Assessment. IEEE/ACM Trans. Comput. Biol. Bioinform. 17, 1918–1931 (2020).

35. Abraham, M. J. et al. GROMACS: High performance molecular simulations through multi-level parallelism from laptops to supercomputers. SoftwareX 1-2, 19–25 (2015).

36. Rollins, Z. A., Faller, R. & George, S. C. Using molecular dynamics simulations to interrogate T cell receptor non-equilibrium kinetics. Comput. Struct. Biotechnol. J. 20, 2124–2133 (2022).

37. Tomasiak, L., Karch, R. & Schreiner, W. Long-Term Molecular Dynamics Simulations Reveal Flexibility Properties of a Free and TCR-Bound pMHC-I System. in 2020 IEEE International Conference on Bioinformatics and Biomedicine (BIBM) 1295–1302 (2020). doi:10.1109/BIBM49941.2020.9313545.

38. Gowers, R. J. et al. MDAnalysis: A Python Package for the Rapid Analysis of Molecular Dynamics Simulations. SciPy 2016 10.25080/Majora-629e541a-00e (2016) doi:10.25080/Majora-629e541a-00e.

39. Valdés-Tresanco, M. S., Valdés-Tresanco, M. E., Valiente, P. A. & Moreno, E. gmx_MMPBSA: A New Tool to Perform End-State Free Energy Calculations with GROMACS. J. Chem. Theory Comput. 17, 6281–6291 (2021).

40. Humphrey, W., Dalke, A. & Schulten, K. VMD: visual molecular dynamics. J. Mol. Graph. 14, 33–38, 27–28 (1996).

41. Springer, I., Tickotsky, N. & Louzoun, Y. Contribution of T Cell Receptor Alpha and Beta CDR3, MHC Typing, V and J Genes to Peptide Binding Prediction. Front. Immunol. 12, (2021).

42. Pettmann, J. et al. The discriminatory power of the T cell receptor. eLife 10, e67092 (2021).

43. Choi, H.-K. et al. Catch bond models may explain how force amplifies TCR signaling and antigen discrimination. Nat. Commun. 14, 2616 (2023).

44. Vita, R. et al. The Immune Epitope Database (IEDB): 2024 update. Nucleic Acids Res. 53, D436–D443 (2024).

45. Springer, I., Besser, H., Tickotsky-Moskovitz, N., Dvorkin, S. & Louzoun, Y. Prediction of Specific TCR-Peptide Binding From Large Dictionaries of TCR-Peptide Pairs. Front. Immunol. 11, (2020).

46. Dieckhaus, H., Brocidiacono, M., Randolph, N. Z. & Kuhlman, B. Transfer learning to leverage larger datasets for improved prediction of protein stability changes. Proc. Natl. Acad. Sci. 121, e2314853121 (2024).

47. Basu, S. & Wallner, B. DockQ: A Quality Measure for Protein-Protein Docking Models. PLOS ONE 11, e0161879 (2016).

