## Supplemental Figure for "AI predicted TCR-pMHC structures differentiate immune interactions"

**Supplementary Methods**

Structural feature extraction

The corresponding contact map of each PDB structure was derived from the per residue alpha carbon (Cα) position. Sites of intermolecular residue interaction were predicted using the findPeaks function from the *pracma* R library (v1.9.9) which is a local maxima peak calling method. Buried surface area (BSA) was calculated from all samples by determining the difference in TCR and pMHC solvent accessible surface area (SASA). Center of mass (COM) calculations was performed based on previous examples^33^, and used to calculate the horizontal displacement distance (r), angle of tilt (ϑ), and angle of twist (φ) of the TCR over the pMHC. The center of mass was calculated for sub molecular regions by the average position of all Cα atoms for each atom in region. Angles between hypervariable regions were determined by a line fit through the region’s extents.

*Folding performance measurements*

When ground truth structures were available, we measured performance using Root Mean Square Deviation (RMSD), Local Distance Difference Test (lDDT), and DockQ. RMSD was calculated as the square root of the distance between the ground truth and predicted backbone position per residue Cα and averaged over all residues. The lDDT is a relative measurement of folding quality that is rotation and translation invariant^34^. The per residue lDDT is the mean of the fraction of preserved atoms within distances of {0.5,1, 2, and 4} angstroms in the predicted structure. We then average global lDDT and regional lDDT for each structure. AlphaFold2 also reports a predicted LDDT (plDDT) value per residue in the b-factor of the output PDB file. While RMSD and lDDT cannot be applied to structures that do not have ground truth x-ray structures, we do calculate the lDDT of predicted pMHC in experimental structures that have a matching pMHC in ground truth structures. DockQ is a global and local evaluation of multimeric structure prediction^35^. Global DockQ score determined by highest chain pair local DockQ score and multimeric quality determined based on previously determined classification (Incorrect = (F_nat_ < 0.1 or (LRMS > 10 and iRMS > 4.0)), Acceptable = (F_nat_ ≥ 0.1 and F_nat_ < 0.3) and (LRMS ≤ 10.0 or iRMS ≤ 4.0) or (F_nat_ ≥ 0.3 and LRMS > 5.0 and iRMS > 2.0)), Medium ((F_nat_ ≥ 0.3 and F_nat_ < 0.5) and (LRMS ≤ 5.0 or iRMS ≤ 2.0) or (F_nat_ ≥ 0.5 and LRMS > 1.0 and iRMS > 1.0)), or High (F_nat_ ≥ 0.5 and (LRMS ≤ 1.0 or iRMS ≤ 1.0)). Where F_nat_ is the proportion of preserved atoms in the binding pocket still within 5 A, Ligand Root Mean Square deviation (LRMS) and interface Root Mean Square Deviation (iRMS) is to deviation measures of the superimposed chains proposed by the CAPRI community^36^.

*Physics calculations*

High energy atomic clashes were calculated using the standard radii for C, O, N, and S^37^. Hydrogen bonds were determined in GROMACS by solvating the TCR-pMHC in water using a charm27 forcefield. Once hydrogen bonds were determined using a bond length cutoff of 2-4 Å and a bond angle >120°, we classified the relative bond strength as “weak” (<120°, 3.2-4 Å), “moderate” (120-170°, 2.5-3.2 Å) or “strong” (>170°, <2.5 Å). Binding energy was calculated using the Rosetta energy scoring function as previously described^38^, on native predicted and relaxed structures. Because we did not observe a linear relationship between calculated structures and experimentally determined binding energies (Extended Data Fig 1i), we also calculated binding energy with prodigy^39^ and PBEE^40^ which use learning models to estimate binding energy calculated from more accurate methods, like Absolute Binding Free Energy (ABFE).

**Supplementary Tables Legends**

**Supplementary Table 1.** Ground truth structure meta data. All structures were downloaded as single pdb files in raw format.

**Supplementary Table 2.** Experimentally validated structure dataset used to predict structures.

**Supplementary Table 3.** Fake structures generated from ground truth and experimental. Base refers to information from the structure that the MHC was derived from and target refers to the information from the structure that the TCR was derived from.

**Supplementary Table 4.** Parameters used in deep learning models that were trained in this study. Dataset for NetTCR2.0 was smaller due to only accepting 9 AA epitopes.

**Supplementary Figures**

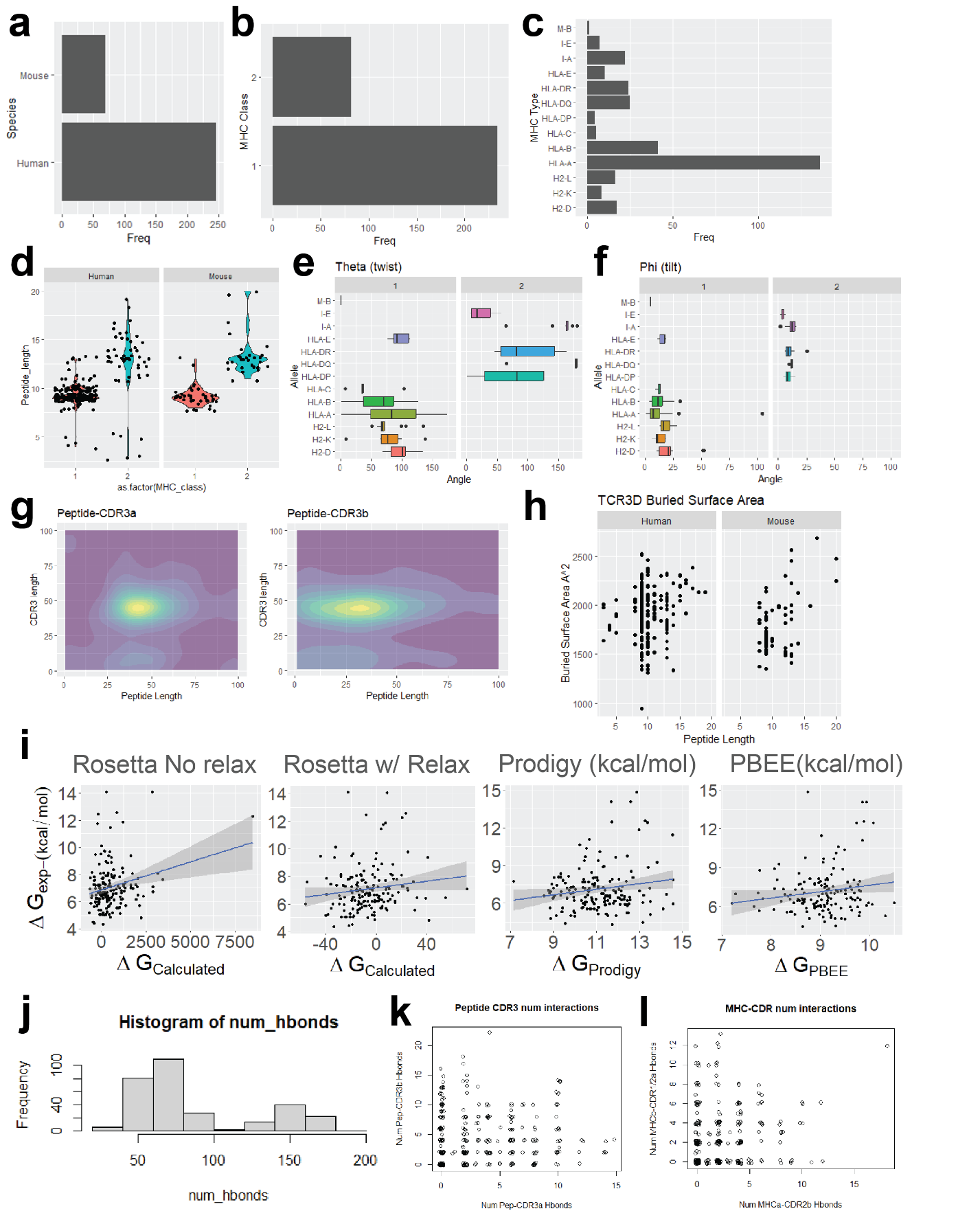

**Supplementary Figure 1.** Ground truth TCR-pMHC structure characteristics were analyzed for properties that could be used as metrics of structure prediction quality and interaction features. **(a-c)** Structures were predominantly from humans and MHC-I interacting TCR structures. There was over-representation of HLA-A, specifically HLA-A2 bound TCR’s due to the overabundance of previous research of HLA-A2 bound antigens. **(d)** Peptide lengths in both human and mouse were on average 9 aa and 13 aa for MHCI- and MHCII-bound structures, respectively. **(e-f)** COM calculated values for the TCR twist (theta) and tilt (phi) over where it sits on the binding pocket of the pMHC molecule. **(g)** Structures had predominantly a single peak at which the CDR3α/β potentially interacted with the peptide which was focused in the middle of the CDR3 region based on a density map of all peaks. **(h)** Buried surface area (BSA) showed a linear relationship with peptide length, showing even coverage by interacting TCRs. **(i)** Real world calculated binding energies (derived from K_d_ measurements) were compared to multiple methods of calculation. Rosetta scoring function was used either with or without full atom relaxation prior to measurement. Prodigy and PBEE were both run using structures as input. Native structures show consistent hydrogen bonding **(j)** structure wide, **(k)** between peptide-CDR3, and **(l)** MHC arm-CDR2/1 residues. Less than 30 structures did not have either a CDR3α or CDR3β hydrogen bond with a median of 8 bonds per structure.

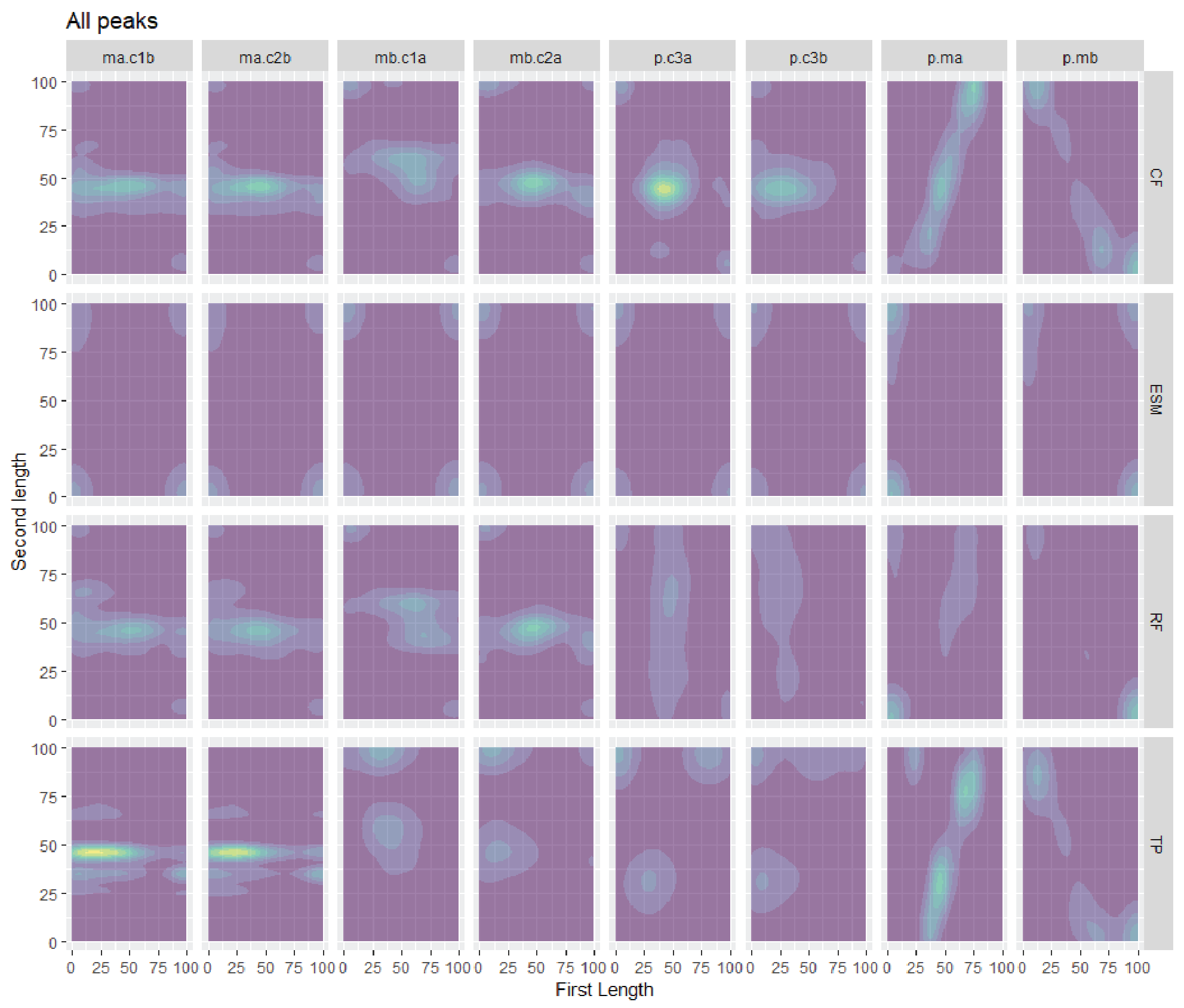

**Supplementary Figure 2.** Density map of peaks for each model by region.

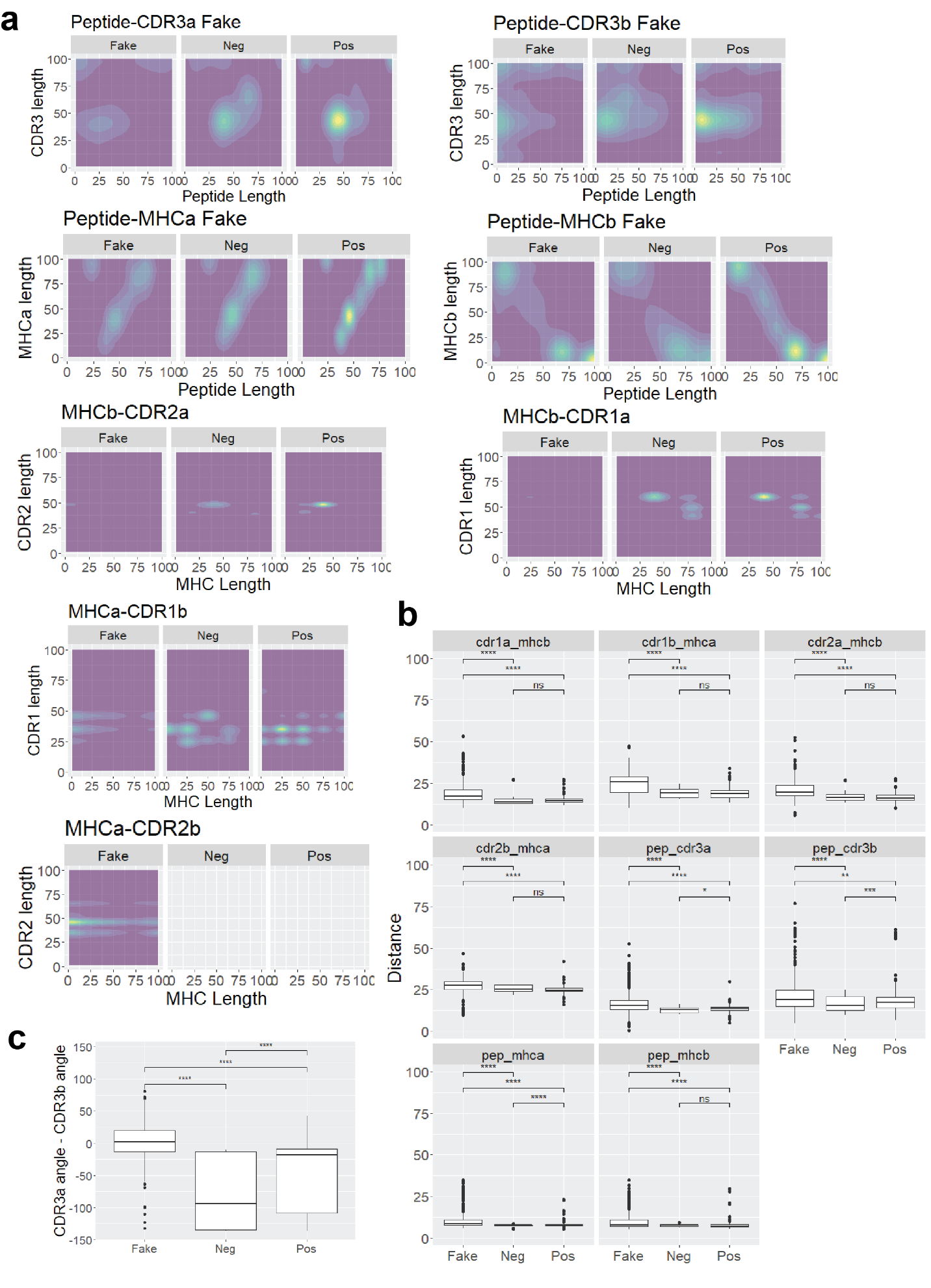

**Supplementary Figure 3.** Characteristics of interacting regions of the TCR and pMHC. **(a)** 2D density plots showing the peak interaction locations of the two regions on the x and y axis. Values 0-100 represent the normalized distance (n-terminal to c-terminal) of each region. Green colors represent areas where more structures were found to have interacting residues as defined by the closest inter-residue distance at least closer than 10 Å. **(b)** Distance between the center of mass of each interacting region. **(c)** The difference between the CDR3α and the CDR3β angle shows that both regions are perturbed in the prediction of a non-interacting structure.

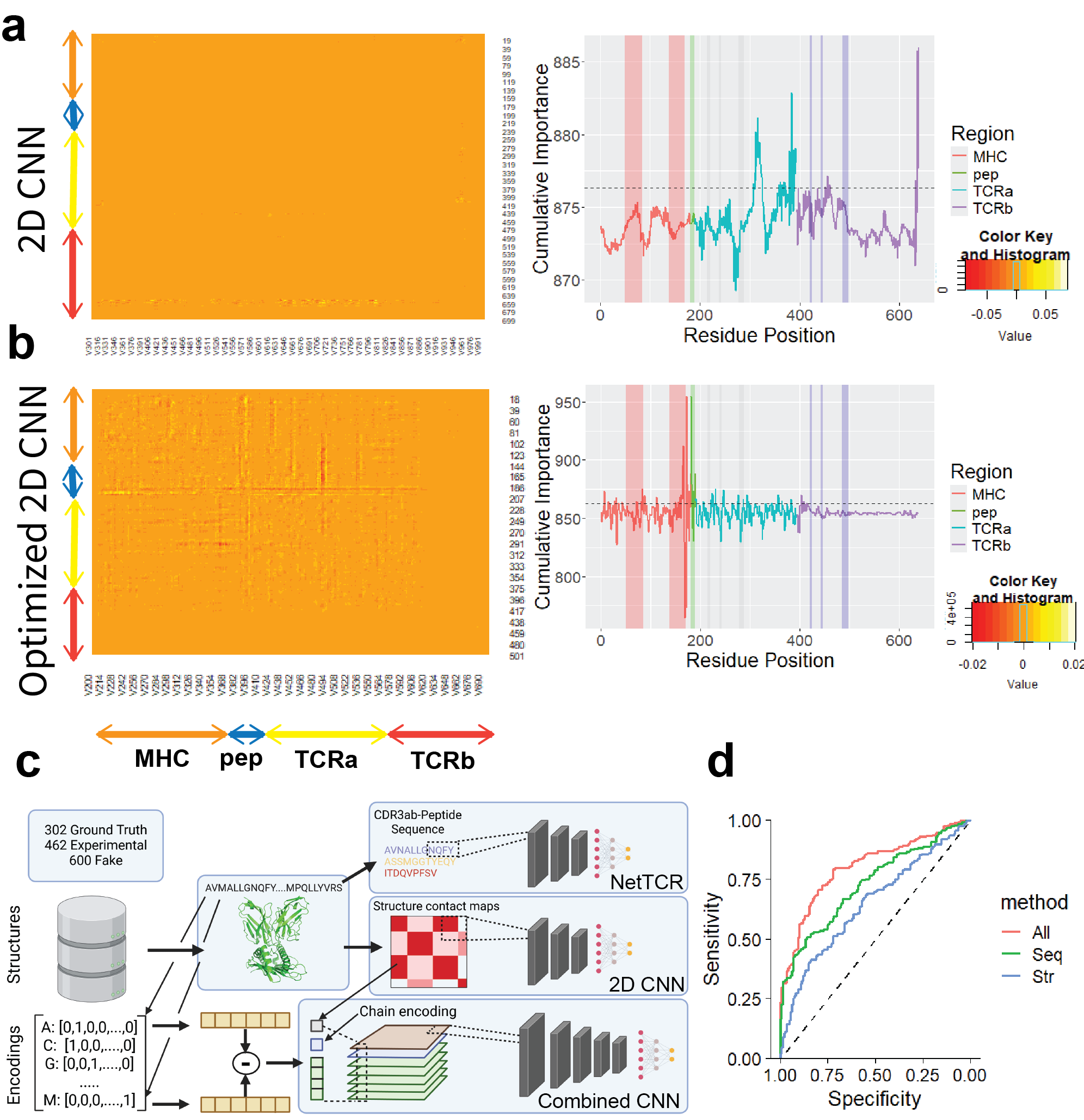

**Supplementary Figure 4.** Deep learning models predict interactions between structures using contact maps. Summed feature importance of test structures is reported based on approximate chain position for **(a)** 2D CNN and **(b)** optimized 2D CNN. Cumulative feature importance at each residue position is reported on the y axis with the horizontal line plotted at the 90^th^ percentile of importance scores. Interacting regions in MHC, peptide, and TCRαβ (CDR1-3) are highlighted by vertical lines. **(c)** General strategy for developing deep learning models is reported. Sequence based features were derived from peptides for NetTCR2.0 but for the combined sequence we used Blosum64 encodings for each amino acid, using the difference between vectors for each pair. Chain encodings were one hot encoded for each chain pair and the physical distance between residue Cα’s was appended to the end of the resultant vector. **(d)** ROC of the combined CNN by which features were included. Black dashed line representing the 50% AUC.

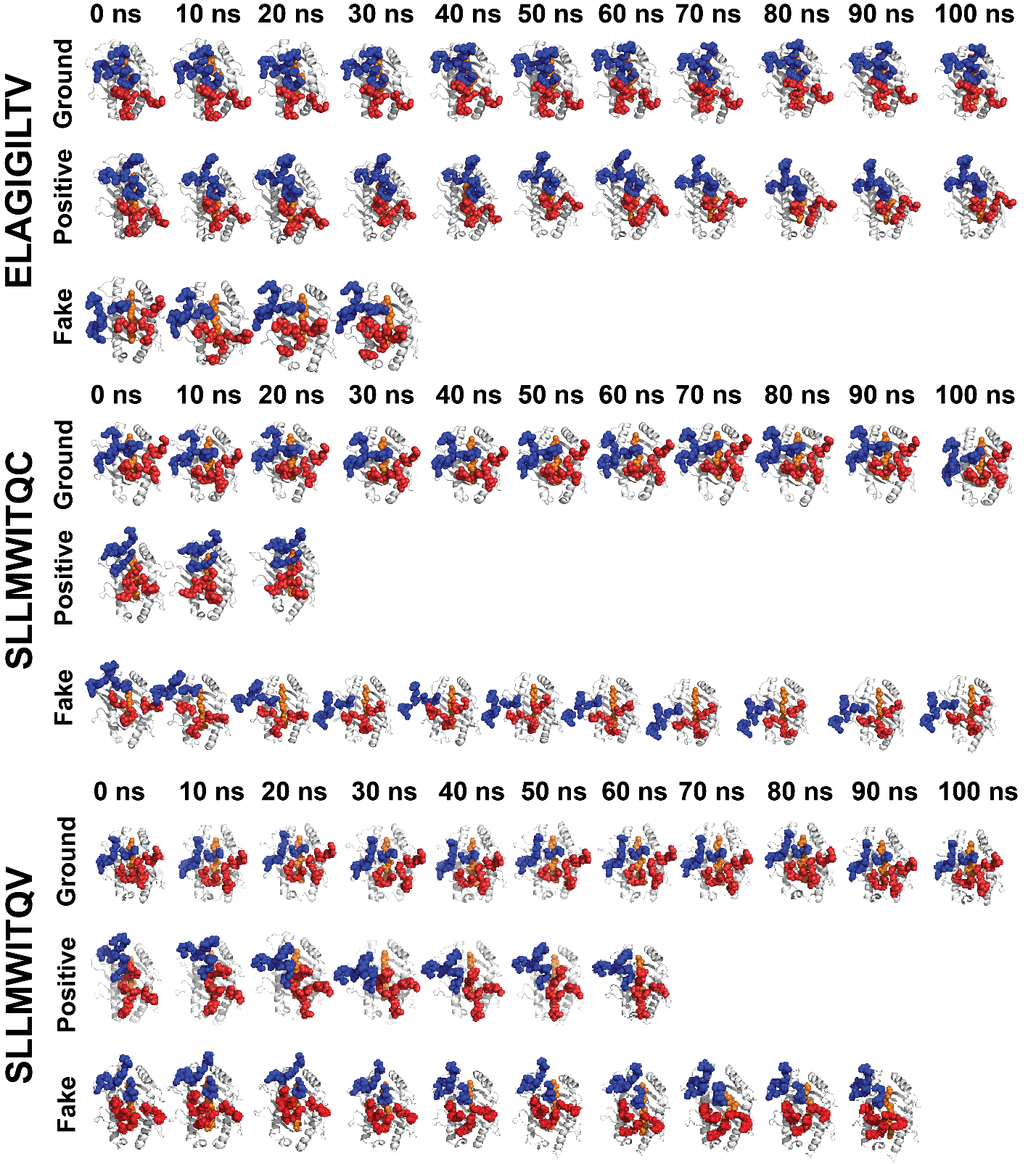

**Supplementary Figure 5.** Examples of simulations run for three peptides that included ground truth, positive and fake structures. Simulations ran up to 100 ns with early stopping for LINCS errors > 1000. PDB structures for ELAGIGILTV (5nht, pos170, fake149), SLLMWITQV (2bnq, pos104, fake190) and SLLMWITQC (2bnr, pos47, fake93) were first solvated with water using a TIP3 model and NaCl ions before equilibration. All simulations were run using an amber99 forcefield in GROMACS software.
